# Kontext: Kernel-based method bridging spatial domain identification and cell-cell communication

**DOI:** 10.64898/2026.10.26.761813

**Authors:** Anthony Ozier-Lafontaine, Nicolas Dray, Paul Villoutreix, Laura Cantini

## Abstract

The spatial organization of biological tissues is characterized by spatial domains: regions with coherent transcriptional and anatomical properties, heavily influenced by local cell-cell interactions. Although identifying spatial domains is a common task in spatial transcriptomics, most existing methods provide limited mechanistic insight into their organization. We present Kontext, a kernel-based framework that jointly represents cells and their local environments, enabling the identification of spatial domains grounded in their underlying ligand-receptor interactions. This representation further provides complementary outputs for characterizing cellular microenvironments, spatial niches, and sender-receiver communication patterns. Kontext outperforms state-of-the-art methods in recovering reference domain annotations across several spatial transcriptomics technologies and biological models, while offering improved computational efficiency and scalability. Applying Kontext on a Xenium temporal mouse heart cryoinjury dataset, we characterize the spatiotemporal evolution of the injury and identify an undescribed sustained activity of the complement system in the communication between fibroblasts and macrophages during scar maturation. In a million-cell three-dimensional Open-ST dataset of human metastatic lymph node tissue, we combine spatial domain identification and niche analysis to define the boundary region between the tumor and its microenvironment, revealing distinct communication patterns between this interface and the tumor core.

## 1 Introduction

Single-cell-resolution spatial transcriptomics technologies provide unprecedented opportunities to quantify gene expression within its native spatial context. Both imaging-based ^1–5^ and sequencing-based techniques ^6,7^ are now capable of measuring hundreds of thousand to millions of spatially resolved expression profiles, while three-dimensional technologies are emerging ^8,9^. These technologies allow to measure a new spatial dimension of cellular identity and state: cells are not only defined by their intrinsic molecular programs but are also shaped by interactions with neighboring cells ^10^. The emergence of spatial transcriptomics data opens the possibility to address new classes of biological questions, including how tissue architecture is established, how gene expression varies across space, how intercellular communication is organized, and how local cellular niches shape transcriptional programs.

The question of tissue spatial structures has primarily been addressed with spatial domain identification methods, which typically apply clustering algorithms to spatially informed embeddings of gene expression. Spatial domain identification methods can be classified in three main families ^11^, statistical models ^12,13^, bayesian models ^14,15^ and different graph-based deep learning methods^16–20^. BANKSY^12^ distinguishes itself through a concatenated embedding that combines expression profiles, with both symmetric and asymmetric aggregations of neighboring expressions. This approach demonstrates that interpretable models remain competitive for spatial domain identification ^11^.

Most existing methods, however, provide limited insight into the biological signals underlying the definition of spatial domains. In particular, ligand-receptor interactions are thought to play a crucial role in shaping and maintaining spatial tissue organization ^21,22^, but methods explicitly linking spatial domains to communication are rare ^23^. Moreover, no gold standard exists for defining spatial domains, ground-truth annotations are still subject to expert biases and largely vary with respect to the field of application ^24^. For instance, brain spatial domains describe tissues and anatomy, whereas embryo spatial domains can describe organs ^6,25^. Except for BANKSY, it is not yet standard practice for most methods to enable adaptation to the various definitions of spatial domains. Finally, as spatial transcriptomic technologies continue to evolve, datasets now routinely comprise tissue slices with hundreds of thousands of cells. However, most current spatial domain identification methods, including deep-learning methods, were not designed for this order of magnitude and have limited scalability ^24^.

We introduce Kontext, a kernel-based framework built on a dual characterization of the cells based on both their expression and local environment to identify spatial domains and their underlying mechanistic origin in terms of ligand-receptor interactions. Kontext relies on the computation of the *“local environment embedding”* which summarizes each cell’s local context as a weighted average of neighboring gene expression profiles, with weights jointly capturing spatial proximity and transcriptomic similarity both encoded with kernels. This dual characterization enables the identification of biologically meaningful spatial domains and provides a basis for interpreting their underlying ligand-receptor interactions. Specifically, Kontext derives cell-cell communication scores as products of receptor expression and local ligand exposure, thereby directly linking spatial domain identification to potential communication mechanisms. The framework further supports a variety of downstream analyses, including local environment influence and niche structure. To address the challenge of spatial domain definition, Kontext allows spatial domains to vary continuously from cell type resolution to tissue-level organization via a tunable parameter that balances the influence of local environment against gene expression. Kontext’s implementation exploits sparse matrix operations to scale to million-cell datasets in minutes without GPU resources.

We systematically benchmark Kontext against state-of-the-art spatial domain identification methods across 30 slices spanning four biological systems and two spatial transcriptomics technologies (Stereo-seq and MERFISH), demonstrating superior domain identification at substantially lower computational cost and memory usage. We then apply Kontext to a multi-sample Xenium dataset of mouse cardiac cryoinjury ^26^, where it reveals temporally structured fibroblast-macrophage communication dynamics at the injury interface. On a three-dimensional Open-ST atlas of a metastatic lymph node ^9^, Kontext distinguishes three regions, the tumor core, its surrounding microenvironment, and the boundary between them, and further characterizes the spatial organization of ligand-receptor interactions between cancer associated fibroblasts and tumor cells.

## 2 Results

### 2.1 Kontext : spatially augmented embeddings for spatial transcriptomic analysis

Cell states are not fully captured by transcriptome alone, they are also shaped by interactions with neighboring cells. Kontext formalizes this duality by associating a cell to two vectors in the same gene space, its expression *X*_*i*_ and its local environment embedding *C*_*i*_, from which spatial domain identification and downstream interpretations in terms of communication, niche composition and local alignment can be inferred (Fig. 1).

**Figure 1:**
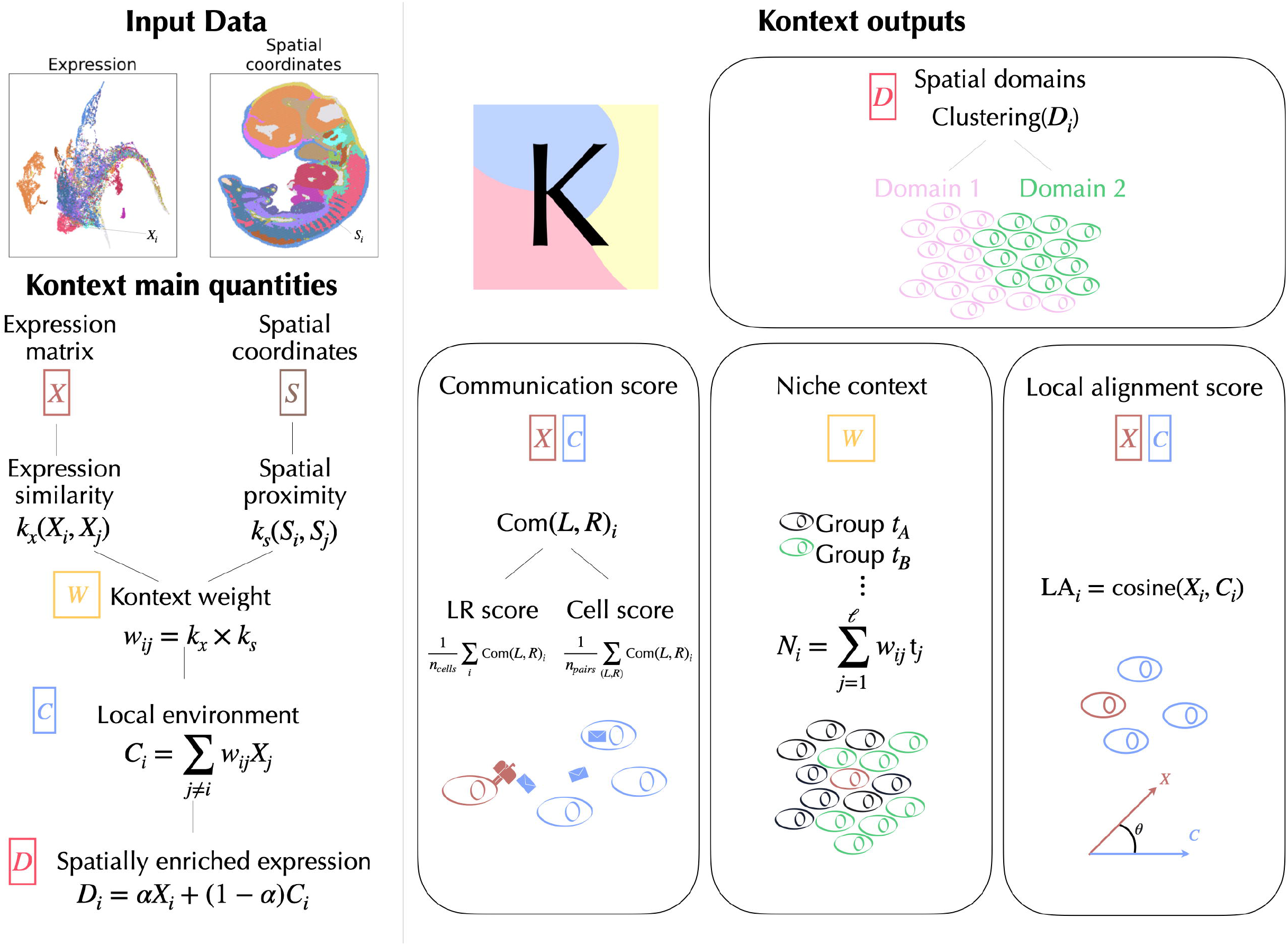
Kontext, a framework bridging spatial domains identification and cell-cell communication. Input data are an expression matrix *X* and associated spatial coordinates *S*. Expression similarity (*k*_*x*_) and spatial proximity (*k*_*s*_) are kernel functions used to obtain Kontext weights *W* from which the local environment embedding *C* and the spatially enriched expression embedding *D* are derived. Spatial domains are obtained from applying a clustering algorithm to*D*. Additional combinations of *X, C* and *W* enable downstream interpretations such as communication, niche and local alignment analyses. Communication score of ligand *L* and receptor *R* in cell *i*, Com(*L, R*)_*i*_, is defined in equation 1. Group indicators *t*_*A*_, *t*_*B*_ and *t*_*j*_ are one-hot encoder vectors of size equal to the number of considered groups.

Consider a normalized spatial transcriptomics dataset of *n* cells and *G* genes (see Methods for data normalization). Cell’s *i* spatial position is *S*_*i*_ in ℝ^*d*^, *d* = 2 for two-dimensional sections and *d* = 3 for three-dimensional volumes. For each cell *i*, Kontext computes a *local environment embedding C*_*i*_ ∈ ℝ^*G*^, whose (g)-th component summarizes the expression of gene *g* in the cell’s local environment. Specifically, *C*_*i*_ is defined as a weighted sum of the expression profiles of its *l* nearest spatial neighbors:

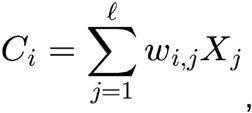

where *w*_*i,j*_ quantifies the contribution of neighboring cell *j* to the local environment embedding of cell *i* . This weight jointly captures the spatial proximity and transcriptomic similarity between the two cells, such that:

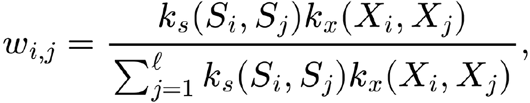

Where *k*_*s*_ is a self-tuned Gaussian kernel on spatial locations defined with respect to *l*^27^ and *k*_*x*_ is a Gaussian kernel on the expression parameterized with the bandwidth size *σ*_*e*_, encoding the range of expression vectors to be considered as similar ^28^. Our default value is *σ*_*e*_ = 1, it corresponds to setting the bandwidth as the median of the distances computed between all cells and their *l* nearest spatial neighbors. Using median distances is standard in kernel methods ^29^ and allows us to focus on local regions in expression space and discard expression vectors too far away. The advantages of coupling spatial proximity with expression similarity is to pull away cells with dissimilar profiles. This approach, inspired from bilateral filters in image analysis^30^, allows sharper local environment transitions at domain boundaries and better detection of distinct populations that are superposed in space. By tuning the parameter *l*, one can control the spatial range of cellular influence in a given tissue.

Our benchmark suggests that *l* = 20, capturing the cell’s immediate environment, provides a good compromise across general settings. This value of *l* = 20 is also adapted to the communication as it fits with the effective intercellular communication distances of approximately 100*µ*m ^31^.

Once the normalized gene expression profile *X*_*i*_ and the local environment embedding *C*_*i*_ are computed for cell *i*, Kontext constructs a spatially enriched expression vector *D*_*i*_ as a convex combination of the two:

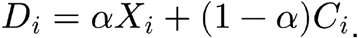

Thus *D*_*i*_ summarizes the expression of a gene either inside the cell or nearby in the tissue, characterizing cells in their local environments. Spatial domains are then identified by clustering these embeddings with a Gaussian mixture model^32^.

One particular feature of Kontext is the possibility to adapt the contribution of expression and local environment in the definition of the spatial domains through the choice of the parameter *α* ∈ [0,1], which is the analogue of BANKSY’s mixing parameter *λ*^12^. Setting *α* = 1/2 corresponds to the absence of *a priori* about the granularity of spatial domains, where a cell and its local environment contribute equally to *D*_*i*_. We suggest this value of *α* as default given its consistent performances in all the benchmark datasets (Supp. Figure S1). Choosing *α* = 0 leads to *D*_*i*_ = *C*_*i*_, corresponding to spatial domains defined as cells facing similar local environments across the tissue, independently of their own expression. Choosing *α* = 1 leads to *D*_*i*_ = *X*_*i*_ which is the input of classical single-cell clustering. Thus increasing from to corresponds to giving more importance to the expression and less importance to the local environment. The possibility to continuously tune the parameter addresses the practical issue of adapting spatial domain definitions across models and applications, from anatomical brain structures, embryonic organ delimitations to tumor microenvironmental compositions.

A key innovation of Kontext is that it goes beyond identifying spatial domains to provide insights into the molecular interactions and local cellular environments that may underlie them. The next paragraphs describe three additional quantities available within the Kontext framework to further interpret Kontext’s spatial domains. The *“communication score”* (Com) provides a proxy to derive possible cell-cell communications from gene expression and local environment. The *“niche embedding”* (*N*_*i*_) leverages the weight matrix to describe a cell local environment with respect to a given categorical label, such as cell type or spatial domain. Finally, the *“local alignment score” (*LA) quantifies local homogeneity by computing the similarity between cells and their local environments.

At the cell *i* level, a proxy of the interaction of a ligand *L* and receptor *R* can be revealed by combining the presence of (*L*) in the context embedding with the expression of (*R*) in the cell’s expression vector. We focus on the communication signals that are exchanged between a sender group of cells *S* and a receiver group of cells ℛ. These groups can be defined as domains, cell types, or any other label of interest. For instance, this approach can be used to compare communication signals between two cell types in two different spatial domains. Given a cell in the receiver group, we quantify its interaction with cells from the sender group through *S* the Kontext communication score

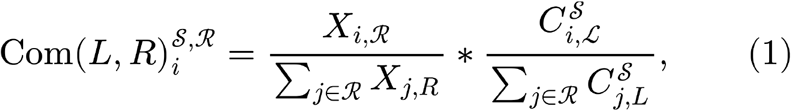

where the scaling with respect to the receiver group makes LR pairs scores comparable to each other (see Methods) and *C*^*S*^ is the contribution of the sender group to the local environment of cell *i*. We obtain a communication score for each cell and each LR pair. This score can be used to derive several quantities : (i) a score per LR pair by averaging over cells and (ii) a score per cell by averaging over LR pairs. Defining the sender group as all the cells, we measure the signals a cell is receiving independently from the sender type. As a consequence, the activity of a LR pair in a subgroup of receiver cells (e.g. cell type or spatial domain) is defined as the mean score of this pair over the subgroup.

Given a set of T cell labels, we use a one-hot encoder to encode each cell label as a vector *t*_*i*_ ∈ ℝ^*T*^. The niche composition of cell *i* can be defined as

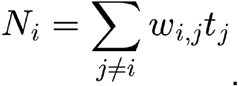

The resulting niche embedding is a vector of size T quantifying the presence of each label around each cell. When the labels correspond to spatial domains, a boundary between two domains can be defined as cells from one domain with a niche embedding of the other domain larger than a given threshold. This niche embedding can also be used in classical niche enrichment analysis.

Finally, local alignment, defined as

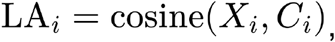

quantifies the concordance of each cell and its local environment and can be interpreted as measures of cellular microenvironment homogeneity. Cells primarily driven by intrinsic programs are more likely to diverge from their surrounding environment, while cells shaped by local context are more likely to align with neighboring cells^33^.

Kontext’s implementation leverages sparse matrix algebra, which makes the computational cost very competitive in terms of time and memory. Kontext is available as an open-source Python package compatible with the scverse ecosystem ^34^, enabling spatial domain identification in a fast, scalable and interpretable way from spatial transcriptomic data (https://github.com/cantinilab/Kontext).

### 2.2 Kontext identifies biologically coherent spatial domains across tissues and technologies

To assess Kontext’s ability to recover spatial domains, we compared its performance against that of three state-of-the-art methods: BANKSY^12^, GraphST^19^ and SEDR^17^. BANKSY was chosen because it relies on a similar approach, GraphST and SEDR are two popular deep-learning methods that have been shown to have competitive performance in a recent benchmark^11^. We compared the methods on MERFISH sections, each assessed at three annotation levels, and Stereo-seq sections, yielding 40 section-annotation combinations spanning four biological systems: mouse brain ^25^, zebrafish ^35^ mouse development ^6^ axolotl brain regeneration ^36^. Sections names, cells and genes counts, and ground truth annotations are summarized in Supp. Tables S1 and S2. We set the number of spatial domains for each benchmarked method to match the author’s original annotation in each section. The MERFISH dataset provides three annotation levels: Allen mouse brain common coordinate framework annotations ^37^, tissue annotations and cell type annotation. Supp. Figure S2 and S3 show the Stereo-seq and MERFISH sections, respectively, colored according to their ground truth annotations.

On MERFISH sections, we leveraged the three levels of annotation to demonstrate the granularity of the recovered domains by varying the value of the parameter *α*. We set for *α* = 0 common coordinate framework regions, *α* = 0.1 for tissue annotations, and *α* = 1 for cell type annotations. For additional information on the impact of on Kontext performance in MERFISH see Methods and Supp. Figure S1. For fair comparisons, we used the same Gaussian mixture model clustering algorithm for all methods, which allows us to fix the target number of spatial domains. Each method was run with different random seeds. As Kontext and BANKSY are deterministic, seeds were used only in the clustering step. For SEDR and GraphST, which involve stochastic embeddings, each seed was used for both embedding generation and clustering. Standard deviations were small compared to the mean scores and comparable across methods, as shown in Supp. Figure S4. We then evaluate the correspondence between the obtained spatial domains and the ground truth cell labels through adjusted Rand index (ARI), normalized mutual information (NMI), homogeneity and completeness as in Hu *et al*.^11^.

Across the 25 Stereo-seq sections, the deep-learning methods SEDR and GraphST failed to process the largest sections containing more than 100, 000 cells. Kontext’s results for these sections are provided in Supp. Figure S5. To ensure a fair comparison, all subsequent benchmarks focus on the 23 sections that could be processed by all methods.

Kontext achieved the highest agreement with the reference annotations on the majority of sections across all metrics considered, obtaining the best score on 25, 29, 32 and 26 sections for the adjusted Rand index (ARI), the normalized mutual information (NMI), homogeneity and completeness respectively (see Fig. 2A and Supp. Figure S5). SEDR ranked second according to ARI and completeness, with the highest score on 7 and 8 sections respectively, while Graph ST ranked second for NMI and homogeneity on 7 and 5 sections respectively. These results were driven largely by zebrafish sections, where SEDR obtained the best ARI on 7 of 12 sections.

**Figure 2:**
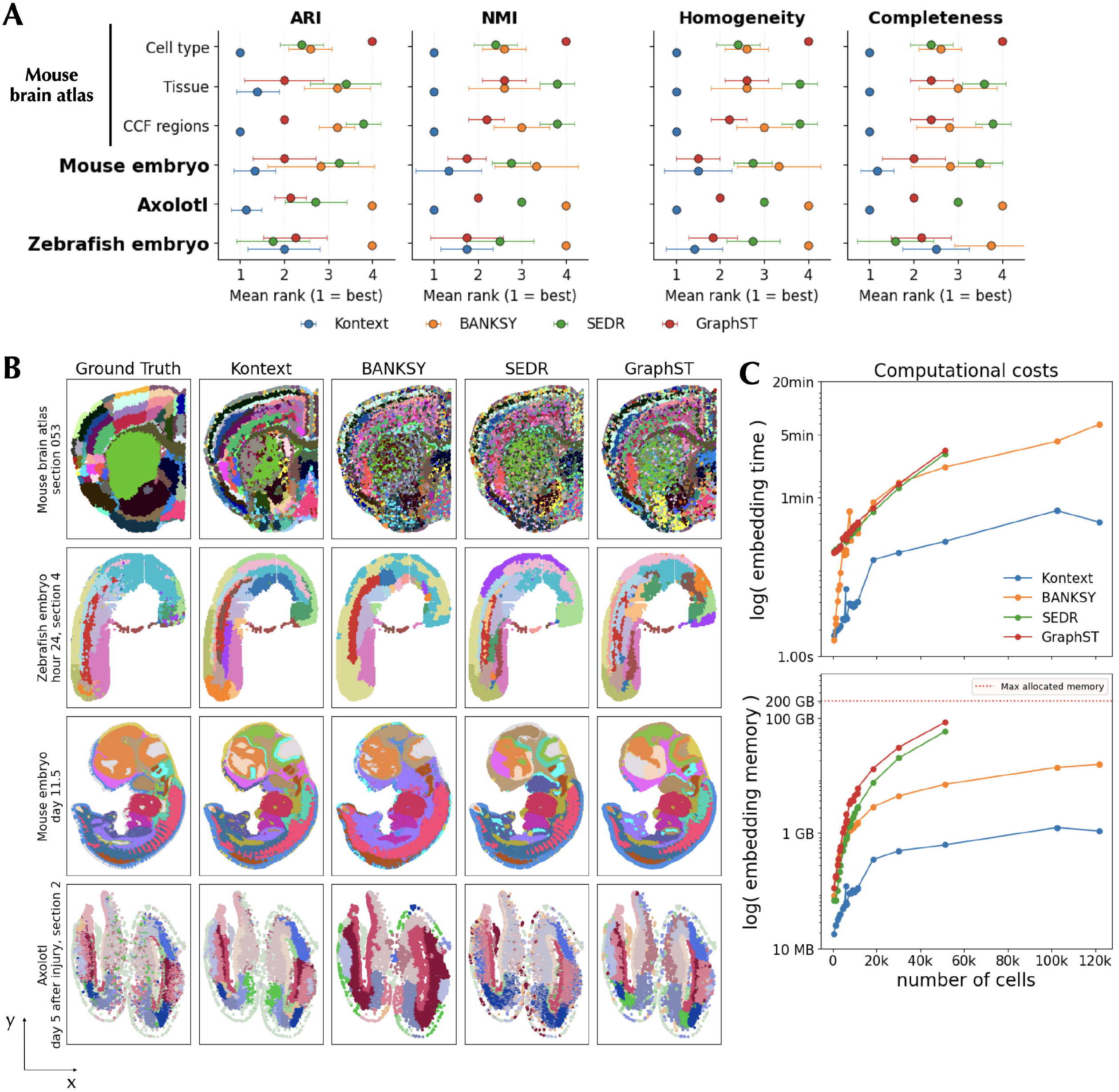
Kontext identifies biologically coherent spatial domains across tissues, technologies and annotations. **A.** Mean rank of each method across the sections of a given dataset, for the adjusted Rand index (ARI), normalised mutual information (NMI), homogeneity and completeness (one panel per metric). Per section ranks were obtained by ordering the four methods with respect to their average metric over 100 random seeds and mean ranks (dots) were obtained from averaging over the sections of each dataset. Horizontal bars indicate standard deviation across sections. Sections on which SEDR and GraphST exceeded the 200 GB memory allocation (MOSTA sections >100,000 cells) are excluded from the ranking. **B**. Representative spatial domain maps for four sections: MERFISH mouse brain atlas (coronal section 53, common-coordinate-framework annotation), Stereo-seq zebrafish embryo (day 24, section 4), mouse embryo (day 11.5) and axolotl telencephalon (day 5 post injury). Left column : expert reference annotation. Remaining columns : domains recovered by Kontext, BANKSY, SEDR and GraphST (seed 0, number of domains fixed to the reference). **C**. Computational cost of the embedding step as a function of the number of cells, across the 25 Stereo-seq sections. wall-clock time (top) and peak resident memory (bottom), both on a logarithmic scale, with points showing the mean over 100 runs for each section. The red dotted line corresponds to the limit of 200 GB memory allocated to each job; SEDR and GraphST (GPU jobs) exceeded this budget for sections above ∼100,000 cells. Non-monotonic variations reflect differences in the number of genes retained per section. Each method is associated with a color, Kontext (blue), BANKSY (orange), SEDR (green) and GraphST (red).

All methods achieved their lowest scores on early zebrafish development stages and their best scores on late-stage mouse embryos and mouse brain sections. This pattern likely reflects both dataset size and the clarity of the ground truth domains: early zebrafish embryo sections are small and lack well defined spatial structures, whereas late-stage mouse embryos and mouse brains are larger and exhibit clearer spatial organization (Supp. Figure S2). Axolotl telencephalon sections feature symmetric structures, with each spatial domain split across two distant regions. Although all methods partially recovered this symmetry, Kontext better matched the reference annotations, suggesting its ability to group cells with similar molecular and environmental information across spatially distant regions (Fig. 2B).

As spatial datasets continue to grow, efficient processing of large cell numbers will become increasingly important. Kontext combines competitive computational efficiency with strong domain recovery, scaling to the largest sections without prohibitive resource requirements or substantial performance loss (Fig. 2C). On the contrary, the deep-learning methods failed to process MOSTA sections exceeding 100,000 cells within the 200 GB memory allocation. Therefore, Kontext not only outperformed other methods in domain detection, but also computed embeddings 12 times faster and with 14 times less memory than BANKSY. Variations in computational costs across sections reflect differences in the number of genes detected.

### 2.3 Kontext resolves the spatiotemporal choreography of macrophage-fibroblast crosstalk during cardiac repair

Cardiac tissue repair is a spatially and temporally orchestrated program in which immune and stromal cells coordinate into successive phases from inflammatory response to scar formation ^38,39^. A central question is whether the spatial organization of this continuous evolution can be described from spatial transcriptomic measurements and further interpreted in terms of Ligand-Receptor programs. Kontext is well suited to this task as it can both identify spatial domains and measure associated communication scores. We employed Kontext to describe the spatio-temporal evolution of eight sections of a Xenium mouse cardiac cryoinjury dataset spanning three time points (sham, 7 days (7D) and 28 days (28D) post-injury) ^26^, the dataset is summarized in Fig. 3A. Using default parameters (*l* = 20, *σ*_*e*_ = 1, *α* = 1/2; Methods), we jointly clustered the concatenated spatially enriched expression embeddings of all sections, spanning sham, 7D and 28D, into eight domains (“SD0-SD7”), each detected in every section (Supp. Figures S6 and S7). The choice of eight domains is aimed at maximizing the signal provided by the injury area (see Methods).

**Figure 3:**
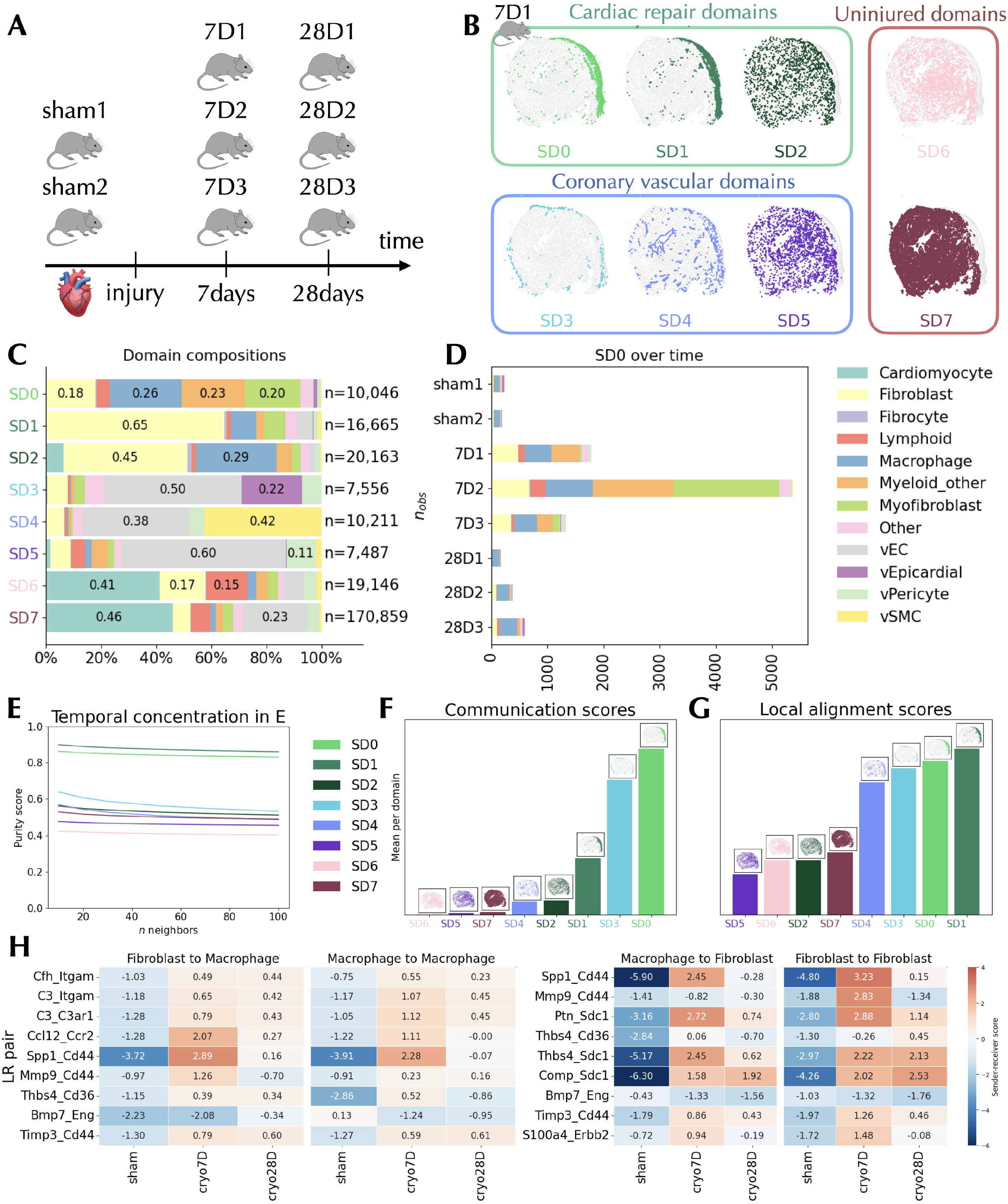
Kontext resolves the spatiotemporal evolution of mouse cardiac reparation. **A.** Description of the dataset, three time points and eight individuals. Sham : sham1, sham2. Seven days post injury : 7D1, 7D2, 7D3. Twenty-eight days post injury : 28D1, 28D2, 28D3. **B**. Spatial localization of the eight Kontext domains on section 7D1. In each panel, cells assigned to one domain are highlighted in the domain color and all remaining cells of the section are shown in grey. Panels are grouped as cardiac repair domains (SD0, SD1 and SD2), coronary vascular domains (SD3, SD4 and SD5) and uninjured domains (SD6 and SD7). **C**. Cell type composition of the Kontext domain as row-normalized stacked bars of the proportion of each main cell type (cardiomyocyte, fibroblast, myofibroblast, macrophage, myeloid other, lymphoid, fibrocyte, vEC, vEpicardial, vPericyte, vSMC, other) within each spatial domain, pooled over the eight sections. The total number of cells per domain is indicated on the right. **D**. Composition of SD0 across sections as stacked horizontal bars of the number of cells of each main cell type assigned to SD0, for each of the eight sections (sham1, sham2, 7D1, 7D2, 7D3, 28D1, 28D2, 28D3). **E**. Temporal purity of the spatially enriched expression embeddings. For each domain, mean fraction of the k-nearest neighbours (k = 10, 20, 30, 50, 70 and 100) sharing the same time point (sham, 7D, 28D), computed in the 50-component PCA of the domain-restricted spatially enriched expression embedding *D*. **F**. Kontext communication scores per domain as mean per-cell Kontext communication score, averaged over all ligand-receptor pairs of the mouse consensus resource, computed for each spatial domain and shown as bars ordered by increasing score. Spatial localization of each domain in section 7D1 is displayed as an indication. **G**. Local alignment scores per domain as mean per-cell local alignment for each spatial domain, ordered by increasing score. Spatial localisation of each domain in section 7D1 is displayed as an indication. **H**. Heatmaps of fibroblast-macrophage ligand-receptor scores across time points. Displayed scores are log2 sender-receiver Kontext communication scores for selected ligand-receptor pairs (rows), aggregated per time point (sham, 7D, 28D; columns). Left panel: macrophage receivers, with fibroblast and macrophage senders. Right panel: fibroblast receivers, with macrophage and fibroblast senders. The colormap is centered in zero, negative values are blue and positive values are red.

The eight spatial domains identified by Kontext can then be readily interpreted based on both cellular composition and spatial localization (Fig. 3B). Domains SD0 and SD1 correspond to the fibrotic niche described by Chan et al.^26^, whereas SD3 and SD4 delineate its exterior and interior borders, respectively. The remaining domains (SD2, SD5, SD6, and SD7) are located in the remote, uninjured region of the tissue (Fig. 3B). Most cells belong to SD7, which represents the healthy tissue domain. SD0 is enriched in macrophages and other myeloid cells, which together account for approximately half of its cellular composition (Fig. 3C). The enrichment of SD0 in myeloid cells, together with the predominance of cells from the 7D time point (Fig. 3C,D), suggests that this domain likely captures the peak of myeloid infiltration observed at 7D in the original study. The domains SD1 and SD2 display similar cellular compositions, consisting primarily of fibroblasts, myofibroblasts, macrophages, and other myeloid cells. Despite these compositional similarities, Kontext distinguishes the two domains based on their distinct spatial contexts: SD1 within the fibrotic niche, whereas SD2 in the remote zone. This illustrates the ability of Kontext to distinguish spatial domains with similar cellular compositions but different tissue locations, capturing spatial context beyond cell-type identity alone.The remaining domains comprise three vascular-associated domains (SD3, SD4, and SD5) located at the exterior border, interior border, and within the tissue, respectively, as well as two domains with a large proportion of cardiomyocytes, SD6 and SD7, predominantly found in the remote zone (Fig. 3B and Fig. 3C).

Focusing on the spatial domains associated with the injury region (SD0-SD1), we observe that SD0 and SD1 exhibit the highest temporal purity across all domains (see Methods), indicating that they are the most temporally distinct ones (Fig. 3E, Supp. Figure S8). This temporal separation is further supported by the UMAP embedding ^40^ of the principal components of each domain (Supp. Figure S9). This suggests that, beyond recovering spatial domains of interest in an unsupervised way, Kontext is able to assign a continuously evolving region observed at different timepoints into one single spatial domain. In addition to their temporal distinctiveness, the injury-associated domains (SD0, SD1) exhibit evidence of stronger communication with their local environment. Indeed, both the local alignment and aggregate communication scores peak in the fibrotic niche domains SD0, SD1. Interestingly, these scores are also high in the vascular domain SD3, possibly explained by the presence of epicardial cells, specific to this domain. More broadly, high local alignment and communication scores characterize injury-associated and vascular domains while low scores correspond to non-injury domains (Fig. 3F and 3G). A high local alignment score can be interpreted as a high concordance between the cells in the spatial domain and their environment, while a high communication score reflects greater ligand-receptor colocalization, suggesting active communication between cells in the spatial domain and their environment. Taken together, these observations suggest that cells in the injury region are intensively communicating and synchronizing their activity, consistent with coordinated cellular responses in injured tissues.

Because several ligand-receptor pairs investigated in the original scRNA-seq study were not represented in the Xenium panel (*Gas6-Axl, Pros1-Axl*, and *Sema3d-Nrp1/Plxna4*), we performed a complementary analysis to assess whether the available Xenium panel could capture biologically relevant cell-cell communication patterns and their temporal dynamics. We focused on the temporal dynamic of the ligand-receptor interactions between fibroblasts and macrophages ^41^. The log Kontext communication scores are shown in (Fig. 3H). Some communication signals were found in one direction only, *Sdc1* and *Erbb* are specific to Fibroblast receivers and *Itgam, C3ar1, Ccr2* are specific to Macrophage receivers only. These pair receptors are known to be specific to these cell types, which stresses the ability of Kontext’s communication scores to catch directed signaling. A complement-associated program involving myeloid receptors *C3ar1* and *CD11b* (Itgam) (*C3– C3ar1, C3–Itgam, Cfh–Itgam*) with macrophage receivers was activated from sham to 7D and remains active at 28D with slightly higher scores when fibroblasts act as senders than when macrophages do^38^. This sustained complement activation at 28D was not reported in the original study. Given that complement signalling is usually associated with inflammation and is expected to subside as tissue repair progresses^38^, its persistence may indicate that macrophages remain engaged in debris clearance during scar maturation (Fig. 3G). Other interactions showed transient activity, peaking at 7D before declining or resolving by 28D. These included *Ccl12*-*Ccr2*, associated with monocyte recruitment^42^, *Spp1*-*Cd44* linked to *Spp1*^*+*^ macrophage activity ^43^, and with fibroblasts as senders, *Mmp9-Cd44* associated with matrix degrading activity and *Ptn-Sdc1* related to growth factor signaling^44^ (Fig. 3H). These temporal patterns are consistent with the reported abundance of *Spp1*^*+*^ macrophages and the monocyte infiltrate at 7D. Interactions involving fibroblasts as receivers also revealed a two-stage matricellular program involving *Thbs4-Cd36* and *Thbs4-Sdc1*. Macrophages were the predominant *Thbs4* senders at 7D, whereas fibroblasts became the main senders at 28D, suggesting a shift towards more autonomous fibroblast activity during scar maturation. Together with the increased activity of *Comp-Sdc1* from 7D to 28D, we interpret this as the activation of myofibroblasts that differentiate into cartilage-like extracellular-matrix during late scar maturation ^45,46^. Finally, we identified additional biologically relevant interactions, including the macrophage-to-macrophage antifibrotic pair *Bmp7-Eng*, which was active in sham tissue but declined at 7D and 28D during fibrosis^47^, and the protease-inhibition-associated pair *Timp3-Cd44*, which was activated at 7D and remained active at 28D. Overall, Kontext communication scores reveal temporally structured interactions between fibroblasts and macrophages that are consistent with their changing roles during tissue repair, from inflammatory recruitment and matrix remodelling to scar maturation.

### 2.4 Kontext reveals CAF-tumor communication at the 3D interface of a metastatic lymph node

Head and Neck Squamous Cell Carcinoma (HNSCC) frequently metastasizes through the lymphatic system ^48^, resulting in spatially structured regions where the tumor-lymph interface influences the progression of the disease ^49^. Cancer-associated fibroblasts (CAFs) are overrepresented at this interface where they play a major role ^50,51^. Their ability to modify the extracellular matrix facilitate tumor cells migration and support vascular remodeling, in addition to creating immune-excluding stromal barriers and influencing immune activity ^52^. Moreover, bidirectional CAF-Tumor interactions through ligand-receptor communication promote adhesion, survival and migration of tumor cells ^53^. We therefore asked whether Kontext could recover the spatial organization of a metastatic HNSCC lymph node and characterize the spatial organization of CAF-tumor communication.

To address this question, we thus focused on the three-dimensional Open-ST dataset ^9^, which contains the expression of one million cells over 19 serial sections along with their corresponding three-dimensional coordinates. Cells are annotated according to 10 main cell types (Fig. 4A, Supp. Table 4, Fig. 4B Supp. Figure S10 ). The Open-ST dataset volume is organized as such: tumor cells are accompanied by infiltrating plasma IgG cells and form a connected structure in the upper sections (“section 23”). The remaining part of the section is mainly populated with fibroblasts, plasma IgA and IgG cells, T cells and macrophages. As we go to the following sections below, two branches of tumor cells separate from the main structure and are surrounded with plasma IgG cells and macrophages. One of the branches is present until “section 23” and the other one spans the whole volume. The second branch reduces to a group of keratin pearl cells from section 18 to the bottom section (“section 2”) (see Supp. Figure S10).

**Figure 4:**
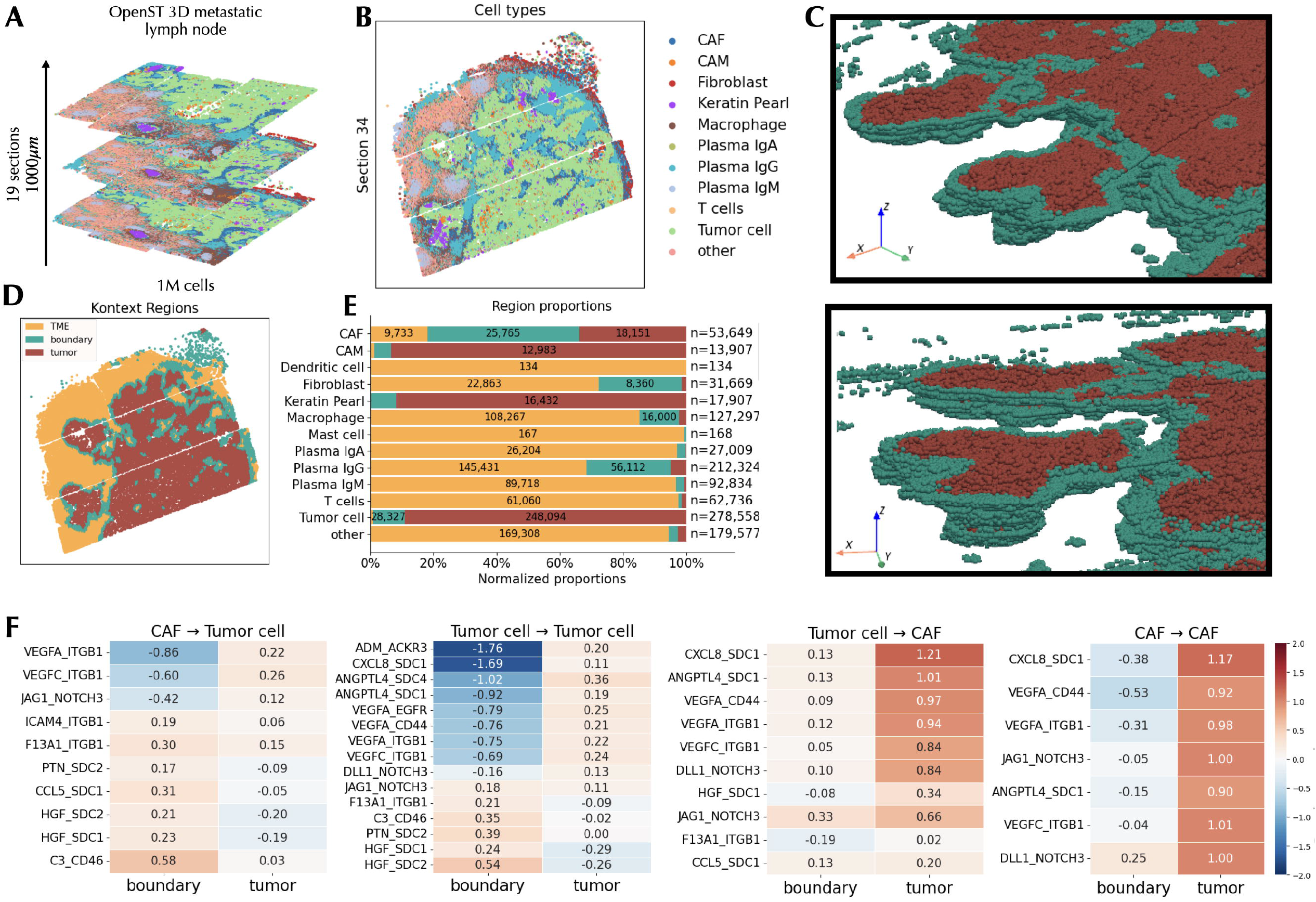
Kontext characterizes the spatial organization of a human metastatic lymph node. **A.** Description of the three-dimensional Open-ST dataset. **B**. Spatial coordinates of the cells of section 34 colored by refined cell type annotation (tumor cell, keratin pearl, CAF, CAM, fibroblast, macrophage, T cell, plasma IgA, plasma IgG, plasma IgM, other). **C**. Two points of view of a three-dimensional point-cloud rendering of the tumor domain and its boundary. Red cells are cells assigned to the tumor domain and teal cells are cells assigned to the boundary region. TME cells are not displayed. **D**. Spatial coordinates of the cells of section 34 colored by regions derived from the Kontext domains and the niche analysis : tumor region, boundary region and TME region. **E**. Distribution of each cell type across the three Kontext regions as row-normalized stacked bars giving, for each cell type, the proportion of its cells located in the TME, boundary and tumor regions, pooled over the 19 sections. The total number of cells per cell type is indicated on the right. **F**. Heatmaps of CAF-tumor ligand-receptor scores across two regions, boundary and tumor. Displayed scores are log2 sender-receiver Kontext communication scores for selected ligand-receptor pairs (rows), aggregated over the boundary and tumor regions (columns). Left panel: tumor cell receivers, with CAF and tumor cell senders. Right panel: CAF receivers, with tumor cell and CAF senders. Pairs are ordered by decreasing tumor-boundary contrast. The colormap is centered in zero, negative values are blue and positive values are red.

We applied Kontext to Open-ST with standard parameters (*l* = 20, *σ*_*e*_ =1, *α* = 1/2; Methods). As we were interested in studying tumor-microenvironment cross talk we set the number of spatial domains to two three-dimensional spatial domains (Supp. Figure S11). Importantly, Kontext natively extends to three-dimensional spatial coordinates thanks to the kernel implementation. To our knowledge, this is one of the first applications of spatial domain identification on a three-dimensional technology such as Open-ST. Moreover, Kontext smoothly scales to this million-cell dataset as the spatial domains were obtained in less than four minutes on a CPU node.

Of the two domains identified by Kontext, the “tumor domain” was enriched in tumor cells, keratin pearls, cancer-associated macrophages (CAM) and fibroblasts (CAF), the other “tumor microenvironment (TME) domain” contained the remaining lymphoid and stromal tissues (Supp. Figure S12). By leveraging spatial environment across serial sections, Kontext recovered the three-dimensional continuity of the tumor domain, including both branches and the keratin pearl region that appears detached from the main tumor in lower sections (Fig. 4C, Supp. Figure S13). In contrast, analysing each section independently produced inconsistent domain patterns, with greater variability in the assignment of boundary cell types, including plasma IgG cells, fibroblasts and CAFs (Supp. Figure S14). This proves the value of taking into account the three-dimensional neighborhood of a cell. Moreover, most infiltrating plasma IgG cells and surrounding macrophages were assigned to the tumor domain in 2D (Supp. Figure S15). These results highlight how Kontext can be readily extended to three-dimensional data, using tissue-wide context to identify coherent spatial domains and structures that may be difficult to resolve through independent two-dimensional analyses. .

We next characterized the invasive margin between the two domains by defining a boundary region comprising tumor-domain cells whose local environment embeddings included contributions from TME cells, and TME-domain cells whose local environment embeddings included contributions from tumor cells. This yielded three interpretable regions: Tumor core, boundary and TME (Fig. 4D). The tumor core and boundary together contained 99% of the tumor cells (89% in tumor core and 10% in boundary region) and 82% of CAFs (34% in tumor core and 48% in boundary region) (see Supp. Figure S16). The boundary also contained plasma IgG cells, fibroblasts and macrophages, consistent with an active stromal and immune interface (Fig. 4E).

To characterize the cell-cell communication signals underlying the spatial organization of the tissue, we analysed the communication scores between CAFs and tumor cells, accounting for differences in sender abundance across regions (Fig. 4F; Methods). Ligand–receptor pairs associated with hypoxia, stress and angiogenesis (*ADM-ACKR3, CXCL8-SDC1, ANGPTL4-SDC1, VEGFA-CD44, VEGFA-ITGB1, VEGFC-ITGB1*) showed higher scores in the tumor core than at the boundary, for both CAF and tumor cell receivers. Because scores were mean-scaled, they cannot be directly compared between receiver types. Nevertheless, this pattern is consistent with a hypoxic tumor microenvironment in which CAFs are exposed and potentially responding to tumor-associated angiogenic and inflammatory signals ^54^. CAFs in the tumor core also express *NOTCH3* once exposed to the ligands DLL1 and JAG1 from both tumor and CAFs, resulting in positive *DLL1-NOTCH3* and *JAG1-NOTCH3* scores, which suggests the presence of myofibroblastic CAFs in the tumor^55^. This pattern is consistent with juxtacrine *NOTCH3* signalling and a potential myofibroblastic CAF phenotype. As *NOTCH3* signalling requires direct cell–cell contact ^56^. In contrast, boundary CAFs to tumor cells signals were associated to complement activity (*C3-CD46*), compatible with local complement regulation ^57^, and hepatocyte growth factors (*HGF-SDC1, HGF-SDC2*) that are known to drive invasion ^58^. Together, these results reveal distinct communication programs across the tumor microenvironment, with core-associated hypoxia, stress, angiogenesis and *NOTCH3* signalling, and boundary-associated complement regulation and growth-related signals.

## 3 Discussion

Conceptually, Kontext builds on the premise that both a cell’s molecular profile and its local environment provide complementary information for spatial transcriptomics analysis. Kontext combines spatial and expression similarities through kernel functions to construct a sparse graph that captures local cellular context. By incorporating ligand-receptor interactions into this representation, Kontext provides a framework for identifying spatial domains whose organization can be interpreted in terms of the underlying cell-cell communication landscape. This interaction-informed view of tissue organization provides the basis for a series of complementary downstream outputs, including local environment embeddings, spatially enriched expression embeddings, communication scores, niche embeddings and local alignment scores.

A key strength of Kontext is that these complementary quantities are derived from a single deterministic *l*-nearest neighbor graph, providing a unified and interpretable framework for spatial transcriptomics analysis. Rather than treating domain identification, environment characterization and cell-cell communication as independent tasks, Kontext connects them within a common representation. Across four biological systems and two spatial transcriptomics technologies, Kontext identified annotated tissue domains more accurately and efficiently than state-of-the-art methods. In particular, it outperformed deep-learning approaches while requiring substantially less memory and computation, even on CPU hardware. Its adjustable balance between spatial proximity and expression similarity also allows domain identification to be adapted to the biological scale of interest. Our applications further demonstrate the value of this framework for interpreting tissue organization and its evolution. In the mouse heart injury dataset, Kontext enabled the analysis of multiple sections across three time points and eight individuals, identifying spatiotemporal domains and characterizing the communication patterns associated with tissue repair. The temporal evolution of communication scores suggested sustained complement-associated signalling during late scar maturation, highlighting how Kontext can generate biologically relevant hypotheses beyond spatial domain identification. In the three-dimensional, million-cell Open-ST dataset, Kontext recovered coherent tumor structures across serial sections and distinguished communication programs associated with the tumor core and invasive margin. These results illustrate how interaction-informed domain identification, combined with local context across sections, can connect tissue architecture to potentially active cell–cell communication and reveal spatial organization that may be difficult to capture through independent two-dimensional analyses.

Kontext nevertheless has several limitations. As with many spatial analysis methods, parameter selection is not fully automated. Although the default parameters performed well across the datasets tested, adjustment remains useful when targeting domains at different spatial and molecular scales. Furthermore, ligand-receptor communication scores inferred from transcript abundance do not necessarily reflect protein abundance, localization or functional activity. Integrating spatial protein measurements could therefore provide a more direct characterization of molecular interactions and strengthen the biological interpretation of the identified domains.

Beyond the applications explored here, the representations derived by Kontext could support additional analyses, including spatially variable gene detection, colocalized gene expression analysis, quantification of the spatial influence of specific cell populations and quality-control tasks such as local post-segmentation denoising. More broadly, as spatial transcriptomics advances towards increasingly large, time-resolved and three-dimensional datasets, Kontext offers a scalable framework for identifying spatial domains grounded in local molecular context and for characterizing the interactions, cellular niches and microenvironmental features associated with them across space, time and tissues.

## Data availability

All datasets used in this work are publicly available. Benchmarked Stereo-seq datasets were obtained from STOMICS database (ZESTA : https://db.cngb.org/stomics/zesta/, MOSTA : https://db.cngb.org/stomics/mosta/, ARTISTA : https://db.cngb.org/stomics/artista/ ), and benchmarked MERFISH datasets were obtained from the cellxgene database (https://cellxgene.cziscience.com/collections/0cca8620-8dee-45d0-aef5-23f032a5cf09). The Xenium spatiotemporal dataset of a mouse heart recovering from an injury and the three-dimensional Open-ST dataset of a metastatic lymph node were loaded from the Gene Expression Omnibus under accession numbers GSE280376 and GSE251926 respectively.

## Code availability

The Kontext python package is available at https://github.com/cantinilab/Kontext. The functions and notebooks to reproduce benchmark results and the analysis of the mouse cardiac injury and metastatic lymph node, as well as to reproduce the figures are available at https://github.com/cantinilab/kontext_reproducibility.

## Acknowledgement

This work was performed using HPC resources from Institut Pasteur. We acknowledge the help of the HPC Core Facility of the Institut Pasteur and Deborah Philipps for the administrative support. We thank Ines Assali for her advice on clustering algorithms.

## Authors contribution

AOL, LC, and PV conceived the study. AOL developed the tool Kontext and performed the computational experiments. ND provided inputs on the methodological decisions. LC supervised the project. AOL and LC wrote the manuscript. PV and ND revised the manuscript. All authors have read, edited, and approved the submission of the manuscript.

## Online Methods

### Notation

We consider *n* cells and *G* genes, described by an expression matrix *X* ∈ ℝ^*n*×*G*^ with rows *X*_*i*_, and by spatial coordinates *S* ∈ ℝ^*n*×*G*^ with rows *S*_*i*_, *d* = 2 for tissue sections *d* = 3 and for three-dimensional volumes. Let denote *N*(*i*) the set of *l* neighboring cells of *i* in *S*, excluding *i*, defined as its *l* nearest spatial neighbors, and *E*_*G*_ = {(*i, j*) : *j* ∈ *N*(*i*)} the *nl* edges of the resulting directed graph.

### Kontext

#### Kernels and context embedding

The weights of a cell are only computed for its *l*^th^ nearest spatial neighbor, which results to a total of *nl* weight computations, with *l* ≪ *n*, instead of *n*^2^ if all weights were computed. Spatial proximity is computed through a self-tuning Gaussian kernel

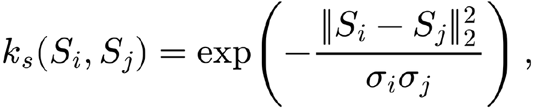

where ∥ · ∥_2_ is the Euclidean distance and 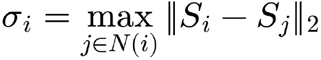 is a cell-specific scaling factor defined as the distance separating cell *i* from its *l*^*th*^ nearest neighbor^27^. Expression similarity is measured with a Gaussian kernel

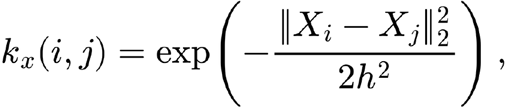

where the bandwidth 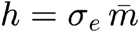 is such that 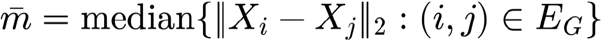 is the median heuristic computed at the edges of the graph and *σ*_*e*_ > 0 is a dimensionless factor. We set *σe* = 1 which corresponds to the median heuristic, the most standard choice for kernel methods in the context of single-cell transcriptomics ^59^. The weight matrix 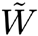, is the asymmetric and zero-diagonal matrix where for pair (*i,j*), we have 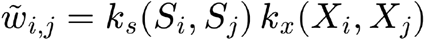. This weight is known in the literature as a kernel based bilateral-filter, a non-linear, edge-preserving smoothing filter used in image processing and computer vision. This choice is particularly adapted to spatial domain detection because it reduces noise while keeping sharp boundaries intact, unlike standard Gaussian filters which blur edges ^30^. Thanks to the filtering on spatial proximity, we have 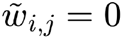 for pairs (*i,j*) that are not connected in *E*_*G*_. Writing 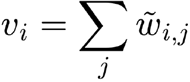 and 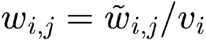, the local environment embedding of cell *i* is 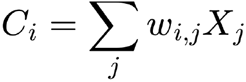.

#### Spatially enriched expression embedding

Given *α* in [0,1],the expression *X*_*i*_ and context embedding *C*_*i*_ are combined into the spatially enriched expression embedding as *D*_*i*_ = *αX*_*i*_ + (1 − *α*)*C*_*i*_

We propose *α* = 1/2 as the default value, which leads to *D*_*i*_ = (*X*_*i*_ + *C*_*i*_)/2. The effect of parameter is discussed below. It is also possible to avoid computing *C*_*i*_ and to compute *D*_*i*_ directly from *X*_*i*_ and 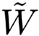 by adding it a diagonal of *αs*_*i*_/(1 − *α*), where *s*_*i*_ is used to scale the expression according to the local environment. The rows of this new matrix sum to 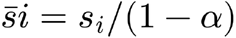. Dividing each row by the value of its sum, we obtain a new weight matrix that has non diagonal values 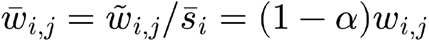, and diagonal values 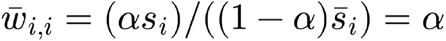.

Thus we have 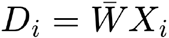.

#### Dimensionality reduction and clustering

The *N*_*D*_ spatial domains are obtained from a Gaussian mixture model on the *p* = 50 principal components of the non-centered spatially enriched expression embedding matrix *D*, computed via randomized truncated singular value decomposition to scale to large datasets ^60^. We run scikit-learn Gaussian mixture model algorithm with ‘reg_covar’ parameter equal to 10^−^3 to stabilize the covariance matrix. The choice of *p* = 50 principal components is a good compromise to distinguish signal from noise in sequencing-based technologies while being not too large for image-based technologies where fewer genes are available. The principal components are computed on the non-centered matrix *D* to preserve its sparsity and enable the method to scale to large datasets.

#### Multi-slice and 3D spatial domain identification

We propose to apply Kontext to multi-slice datasets in two different ways depending on the distance between the sections. For slices far apart from each other, the embeddings are computed separately for each slice. Otherwise on 3D datasets, Kontext can be applied directly on the three-dimensional coordinates of the cells. We applied Kontext to multiple tissue sections in the mouse heart injury dataset. Weights and embeddings were computed on each preprocessed section separately with respect to one shared panel of genes. Based on the assumption that local environments are comparable across sections, dimensionality reduction over the spatially enriched expression embeddings is performed on the concatenated dataset, resulting in one shared low-dimensional space, from which we determine the spatial domains. For 3D data such as the Open-ST metastatic lymph node dataset, we analyse it as a unique dataset, where distances are computed from the three-dimensional coordinates and cells may have neighbors in several consecutive slices.

#### Communication score

In order to take into account the amplitude of the expression of the ligands and receptors in the communication scores, we consider 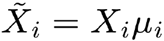 instead of *X*_*i*_, where *µ*_*i*_ is the L2-norm of the cell before the L2-normalization step of the preprocessing (see Preprocessing).

Let *L* and *R* be the ligand and receptor of a ligand-receptor pair. We focus on the possible interactions between ligands *L* derived from a sender group of cells and receptors *R* derived from a receiver group ℛ containing *n*_ℛ_ cells. The sender-specific context of cell *i*, 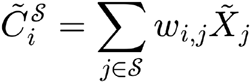 measures the specific contribution of sender group *S* to the local environment of cell *i*.

As a local interaction is only possible when a receiver cell *i* has at least one sender cell in its local environment, we restrict the analysis to receiver cells where 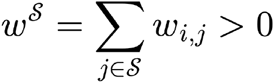. To quantify the contribution of sender cells in the environment of receiver cells independently from their local abundance, we define the normalized sender-specific context 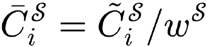. Using 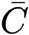 instead of 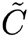 allows to account for the large discrepancies in local sender availability in the different regions of a tissue, in order to focus on differences in ligand availability per sender. As rows are normalized to sum to 1, 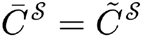 when the sender group is the whole dataset. The communication score of cell *i* from a receiver group ℛ expressing receptor *R* with respect to ligand *L* from sender group *S* is defined with

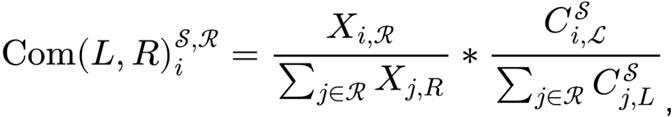

The mean-scaling step is applied to make ligand-receptor pairs comparable with each other within a receiver group and across different sender groups. However, receiver-based mean-scaling makes scores involving different receiver groups non comparable. Throughout this work, we focus on log-communication scores, meaning that a positive score identifies a cell that expresses the receptor *R* more than the average receiver cell in the presence of sender cells distributing more ligands *L* than in the average environment of receiver cells. Ligand-receptor pairs are taken from the organism-matched LIANA+ resource ^61^, restricted to pairs whose two partners are measured. We also filter the pairs where the receptor is expressed in less than *t*_ℛ_% of the receiver cells.

#### Niche composition

For a categorical annotation such as cell-types or spatial domains, encoded into a one-hot indicator *T*_*j*_, the niche embedding of cell *i* is 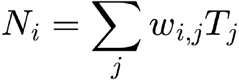, interpreted as the vector of weighted proportions of each category in its environment. Applied to domain labels, *N*_*i*_ also defines interfaces: a cell whose niche contains a non-zero proportion of another domain, represented by *N*_*i,t*_ > 0 is assigned to the boundary between the two, which is the rule used to partition the lymph node into tumor region, boundary region and TME region.

#### Local alignment

The local alignment of cell *i* is the cosine similarity between its own expression and its local environment profile,

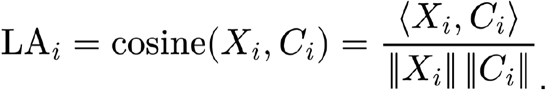

Biologically, cells with high concordance are transcriptionally aligned with their surroundings and thus likely shaped by their local environment, whereas cells with low concordance follow programs that diverge from their neighbors. Averaged over a spatial domain, LA ranks domains by the transcriptional homogeneity of their local environment.

#### Label purity of a spatial domain

Given a categorical label (e.g. timepoint), the purity score is computed within each domain. To do so, the *k* nearest neighbors of every cell of a domain are computed in the PCA space of the embedding, and the purity score for *k* is the average number of its neighbors sharing its label^62^. Purity can be compared across dimensionality-reduced spaces derived from *X, C* and *D* to indicate which representation carries the label-specific signal. A clear separation between different categories results in a high purity.

#### Complexity and implementation

Given a spatial transcriptomics dataset of *n* cells and *l* considered nearest neighbors, all quantities are supported on the *nl* edges of the spatial graph that is sparse. Sparse matrix algebra allows Kontext complexity to scale as *O*(*nl*) to obtain the weight matrix and 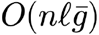 for the local environment embedding, 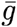 being the mean number of genes detected per cell. Kontext is implemented in Python on AnnData ^63^ and scanpy ^64^. Sparse matrices were handled with scipy ^65^, many standard machine learning tools were imported from scikit-learn ^66^ and array oriented computations were optimized via numba ^67^.

### Datasets

#### Stereo-seq

Twenty-five published sections from three systems, with the original annotations as reference domains: twelve zebrafish embryo sections over six developmental stages (ZESTA^35^; samples zf3, zf5, zf10, zf12, zf18, zf24, one to four slices each). Seven axolotl telencephalon sections from 2 to 60 days post injury (ARTISTA)^36^, for which coordinates were reflected along both axes. Three cells with extreme coordinates were excluded from the section corresponding to 20 days post injury, corresponding to the two cells with minimal horizontal axis coordinate and one cell with maximal vertical axis coordinate. Six mouse embryo sagittal sections from E9.5 to E16.5 (MOSTA, slice E1S1), annotated by organ^6^. Raw counts were used as input throughout.

#### MERFISH

Five coronal sections of the whole mouse brain atlas^25^ (animal 1) were evaluated at three annotation levels provided with the atlas, Allen Mouse Brain Common Coordinate Framework region^37^, tissue and cell type. Each section annotation pair is treated independently. Cells unannotated at the level considered were excluded.

#### Xenium cardiac cryoinjury

The mouse cardiac cryoinjury dataset^26^ contains eight sections over three conditions (sham, 7 and 28 days post cryoinjury; two, three and three sections). Count matrices, feature tables and centroids were read from the deposited Market Matrix files, with the published cell-type labels, except section 7D2 for which the authors’ resegmented annotation was used. The 24 fine-grained cell types provided by the authors were aggregated into 12 main cell types (Supplementary Table 3). Cells without a label were excluded, as were two single cells in sections 28D2 and 28D3 respectively (removed cells were indexed as hmonjnan-1 in section 28D2 and mifmgime-1 in section 28D3)

#### Open-ST metastatic lymph node

The three-dimensional atlas of a Head and Neck Squamous Cell carcinoma metastatic lymph node^9^, composed of 19 registered slices was analysed using the aligned three dimensional coordinates and cell type annotations published by the authors. Cell type annotations were renamed (Supplementary Table 4).

#### Preprocessing

Kontext is a framework within which all the underlying data structures are kept sparse throughout the whole pipeline. This makes Kontext scalable and computationally cheap, but an adapted preprocessing procedure is needed. We propose two technology specific preprocessing, one for sequencing-based technologies and one for imaging-based technologies.

##### Preprocessing for sequencing-based data

For sequencing-based technologies where tens of thousands of genes are measured, we propose to remove genes present in less than 20 cells and to select the 2000 most highly variable genes (Scanpy implementation, Seurat flavour, computed on normalized values), then we apply a cell-wise normalization to 10^4^ with respect to the total counts over all selected genes. We denote by *Z* the filtered and scaled count matrix. The log-normalized expression matrix used as Kontext input is then defined as 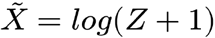.

##### Preprocessing for imaging-based data

For imaging-based technologies, there is no gene filtering nor scaling, the preprocessing reduces to the log-normalization step. Finally, an additional L2-normalization step is applied to 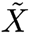, so that 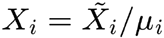, where 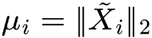 that corresponds to setting all cells as ∥*X*_*i*_∥_2_ = 1. This has three consequences that the model relies on: the squared Euclidean distance between two profiles becomes proportional to their cosine dissimilarity, so that the expression kernel *k*_*x*_ is a cosine similarity kernel independent of sequencing depth, and every neighboring cell contributes with the same magnitude to the weighted average *C*_*i*_, so that the local environment reflects the composition of the environment rather than the depth of its cells. With this step, all cell expression vectors lie in the unit sphere, which makes their weights and local environment embedding more comparable.

### Benchmarking

#### Compared methods

Kontext was compared with BANKSY^12^, SEDR^17^ and GraphST^19^, run with their published implementations and default parameters unless stated. Each method SEDR, GraphST, BANKSY was run with its author-recommended preprocessing, from the same raw counts. BANKSY used library-size normalization and 2,000 highly variable genes; SEDR used gene filtering (at least 50 cells, 10 counts), normalization to 10^6^, 2,000 highly variable genes on counts and gene-wise standardization; GraphST was run on raw counts with its internal preprocessing depending on the sequencing technology.

We used the following parameters. BANKSY:, azimuthal Gabor filter order, scaled Gaussian weighting, principal components, and mixing parameter, selected as the best-performing value over *λ* ∈ {0.1, 0.2, 0.3, 0.4, 0.5, 0.6, 0.7, 0.8, 0.9, 1} across all sections and metrics at seeds each (Supp. Figure S17). The parameter *λ* in BANKSY is equivalent to (1 − *λ*) in Kontext, thus we show the same tendencies for this parameter across the benchmarked dataset than observed with Kontext’s *α*. SEDR: 200 input principal components, 12-nearest-neighbor graph, default architecture, 200 epochs, learning rate 0.01, 10 internal clusters, the gaussian mixture model algorithm was applied on the 28-dimensional output embedding. GraphST: technology-specific preprocessing, 64-dimensional embedding projected onto 50 principal components.

#### Kontext parameters

##### Balance parameter (*α*)

Kontext’s spatial domains rely on the choice of parameter *α*, that represents the importance given to both expression and local environment. We propose *α =* 1/2 as the default value as it corresponds to giving the same importance to both. These values also show competitive empirical performances on the benchmarked datasets (results at fixed *α =* 1/2 are in Supp. Figure S18). However, we showed on the MERFISH sections that choosing the value of *α* based on the ontology of the desired annotation can lead to increased performances, common coordinate framework regions are anatomical regions for which the local environment alone should be sufficient to detect them, tissue annotations are both anatomical and functional structures, that should be recovered from a spatially enriched expression embedding mostly based on local environment, and cell types annotation were defined from expression alone. Here we extensively explore the sensitivity of the spatial domains with respect to *α*. Varying *α* across {0, 0.1, 0.2, 0.3, 0.4, 0.5, 0.6, 0.7, 0.8, 0.9, 1}, the performances aligned with the expected behavior (Supp. Figure S1). Common coordinate framework regions were best detected when *α* was close to 0 and rapidly decreased as *α* increased. Tissues were fairly detected for all values of *α* inferior to 0.4, and agreement between identified spatial domains and cell types increased monotonically as *α* increased to 1. This analysis demonstrates the adaptability of Kontext to fit a large variety of annotation definitions.

##### Other parameters

We set Kontext parameters as l = 20, *σ*_*e*_ = 1, *p* = 50, *α =* 1/2 on all Stereo-seq sections. For MERFISH mouse brain annotations, *α =* 0,0.1, and 10 for Common Coordinate Framework, tissue and cell-type annotations respectively, fixed a priori from the semantics of each level rather than from performance. We explored the effect of parameters *σ*_*e*_ and *L* on the benchmarked sections (Supp. Figures S19 and S20). Results show that varying *σ*_*e*_ within {0.2, 0.4, 0.6, 0.8, 1, 1.2, 1.4,1.6, 1.8, 2} does not have a huge impact on the performances. However, varying the number of considered neighbors *L* within {5, 20, 100, 300, 1000} has an important impact that is heterogeneous across datasets, suggesting that the area of influence of the cells within a tissue in terms of spatial organisation may vary a lot across models and organs. For instance, best scores on axolotl telencephalon and mouse embryo are obtained with small values of *L* within 5 and 20, while setting *L* = 100 makes Kontext better than state of the art methods on zebrafish embryo sections, and setting *L =* 500 drastically increases the already good performances on the two anatomical annotations of the mouse brain atlas dataset (common coordinate framework and tissue). Overall, setting the parameter *L* = 20 seems to be a good compromise to handle general settings. Note that increasing the parameter *L* also has an impact on the communication scores, as more sender cells will be taken into account as sending communication signals, which can make sense for cells known to have large areas of influence such as neurons but can be physically unlikely for other cell types. Also, increasing *L* decreases the sparsity of the weight matrix, resulting in increased memory usage and computational time (Supp. Figure 21).

#### Evaluation

For fair comparisons, we used the same Gaussian mixture model clustering algorithm for all methods, independently from the author’s proposed clustering algorithms, as it allows to fix the target number of spatial domains. All embeddings were partitioned by the same Gaussian mixture implementation with *K* set to the number of categories of the ground-truth annotation, at 100 random seeds per section. Kontext and BANKSY being deterministic, seeds affected clustering only. For SEDR and GraphST each seed was used for both embedding and clustering. Scores were averaged over seeds per section, and methods compared by mean rank per dataset and through the number of sections at which each attained the highest mean score.

#### Evaluation metrics

The metrics used in this benchmark are inspired from the metrics used in a recent benchmark^11^. Agreement between a recovered partition *V* = {*V*_1_, …, *V*_*K*_} and the reference annotation *U* = {*U*_1_, …, *U*_*R*_} of the *n* retained cells was quantified by four external indices computed from the contingency table *n*_*ab*_ = ∣ *U*_*a*_ ∩ *V*_*b*_ ∣, with margins *a*_*a*_ *=* ∣ *U*_*a*_ ∣ and *b*_*b*_ *=* ∣ *V*_*b*_ ∣. The adjusted Rand index compares the number of cell pairs classified consistently by the two partitions, the score is adjusted with respect to a baseline with respect to a random model :

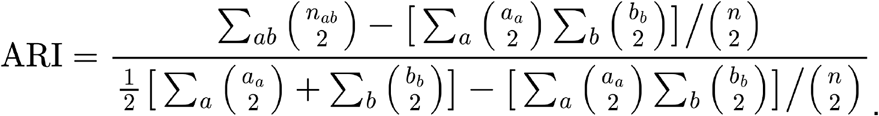

ARI is equal to 1 for identical partitions and to 0 in expectation for independent ones. The three remaining indices are information-theoretic. With

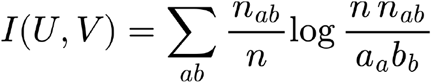

the mutual information and *H*(·) the entropy of a partition, the normalized mutual information, homogeneity and completeness are

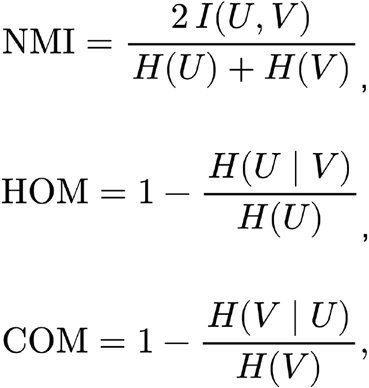

all in [0,1] and equal to for identical partitions. Homogeneity and completeness are complementary metrics : homogeneity decreases when a recovered domain mixes several ground truth categories, completeness decreases when a ground-truth category is split across several recovered domains, so the pair distinguishes over-merging from over-fragmentation at equal ARI. All four were computed with scikit-learn (adjusted_rand_score_,normalized_mutual_info_score,homogeneity_score, completeness_score) on the cells retained after the exclusions described in Datasets, identically for every method. For each section and method, scores were averaged over seeds.

#### Spatial maps colors

Spatial maps were plotted by matching domain colors to the reference labels by maximum-overlap one-to-one assignment (Hungarian algorithm) so that colors are comparable across panels.

#### Runtime and memory

Wall-clock time and memory usage were compared on the embedding step only, excluding loading, preprocessing, clustering and scoring. Peak GPU memory of deep-learning methods was measured from the CUDA allocator. Jobs were run on a SLURM cluster with up to 200GB memory, a job was recorded as failed when it exceeded the memory limit or the time limit of 600min.

### Case-study analyses

#### Mouse cardiac cryoinjury

The eight sections weights and embeddings were computed separately with the same set of parameters and jointly analysed for spatial domains identification and downstream analyses. Kontext was run with *L* = 20, *σ*_*e*_ = 1, *α =* 1/2, the eight spatially enriched expression embeddings were row-concatenated before running a shared principal component analysis and a single Gaussian mixture fit with *N*_*D*_ = 8 spatial domains at seed . This value *N*_*D*_ of was the smallest value resolving the injury-associated compartments while keeping every domain present at every time point. Values of *N*_*D*_ ∈ {2, 3, 4, 5, 6, 7} are shown in Supp. Figure S22. Domains are referred to by index and were annotated from their cell-type composition and their localization relative to the injury. Local alignment and communication scores over all measured pairs were computed per cell and averaged per domain and section. Temporal purity was computed within each domain against the time-point label in *X, C* and *D*, over *p* ∈ {10, 20, 30, 50, 70, 100}. Sender-receveir communication between fibroblast and macrophage cells were computed in both directions, scores were averaged over receiver cells of the relevant type within each section, and pairs were considered if they were expressed in at least 10%of the receiver cells (*t*_ℛ_ = 10).

#### Three-dimensional metastatic lymph node

Kontext was run on the whole 19-slice volume as a single object. The 16 fine-grained cell types provided by the authors were aggregated into 10 main cell types (Supplementary Table 4). During the preprocessing, we added to the 2000 most highly variable genes all the ligand and receptor genes of the organism-matched LIANA+ resource. We used the standard parameters *l* = 20, *σ*_*e*_ = 1, *α* = 1/2 and computed *N*_*D*_ = 2 domains at seed 0. The two domains were labelled tumor domain and tumor microenvironment (TME) domain from their cell-type composition. Three regions were defined from a niche analysis of the two domains. Cells whose niche embedding of the opposite domain was non-zero were assigned to the boundary region, the others to the tumor region or the TME region. This three-way partition is directly derived from Kontext weights and domain labels, thus requiring no additional parameters. Domains and region cell type compositions were computed as contingency tables. In each region, sender-receveir communication between cancer-associated fibroblasts (CAFs) and tumor cells were computed in both directions, scores were averaged over receiver cells of the relevant type within each section, and pairs were considered if they were expressed in at least 15% of the receiver cells (*t*_ℛ_ = 15). The reported genes are the pairs having the highest communication contrast between the two compared regions and they are ordered by decreasing tumor to boundary contrast. The detached tumor subpopulation was identified by visual inspection and used in no quantitative comparison. Point-cloud rendering was performed using the python package PyVista ^68^.

#### Statistics and reproducibility

All random seed values were explicitly stated, from to for the benchmark and for the two case studies. Analyses were run in Python 3.12 with scanpy [≥ 1.9.8], anndata [≥ 0.10.0], scikit-learn [≥ 1.3], scipy [≥ 1.11], numpy [≥ 1.24], numba [≥0.58], liana [≥ 0.1] and leidenalg [≥ 0.9]. Benchmarked packages were banksy-py [≥ 1.3.4] and graphst [≥ 1.1.1]. SEDR was installed from its github repository (https://github.com/JinmiaoChenLab/SEDR.git).

