## Supplementary tables and figures for "Kontext: Kernel-based method bridging spatial domain identification and cell-cell communication"

| | dataset | section | name | n_cells | total genes | n_domains |
|---|---|---|---|---|---|---|
| **0** | ZESTA | zf3_1 | hour 3, section 1 | 1167 | 6261 | 3 |
| **1** | ZESTA | zf5_10 | hour 5, section 10 | 563 | 1962 | 7 |
| **2** | ZESTA | zf10_14 | hour 10, section 14 | 1231 | 2756 | 8 |
| **3** | ZESTA | zf12_8 | hour 12, section 8 | 2081 | 4963 | 11 |
| **4** | ZESTA | zf12_9 | hour 12, section 9 | 2588 | 5802 | 11 |
| **5** | ZESTA | zf18_7 | hour 18, section 7 | 3475 | 7522 | 14 |
| **6** | ZESTA | zf18_8 | hour 18, section 8 | 3048 | 9217 | 14 |
| **7** | ZESTA | zf18_9 | hour 18, section 9 | 5935 | 12485 | 14 |
| **8** | ZESTA | zf24_4 | hour 24, section 4 | 5271 | 12680 | 19 |
| **9** | ZESTA | zf24_7 | hour 24, section 7 | 6318 | 13786 | 18 |
| **10** | ZESTA | zf24_9 | hour 24, section 9 | 4708 | 13042 | 17 |
| **11** | ZESTA | zf24_10 | hour 24, section 10 | 5968 | 13527 | 18 |
| **12** | ARTISTA | 2DPI_1 | day 2 after injury, section 1 | 7668 | 27324 | 18 |
| **13** | ARTISTA | 5DPI_2 | day 5 after injury, section 2 | 8178 | 29442 | 17 |
| **14** | ARTISTA | 10DPI_3 | day 10 after injury, section 3 | 10002 | 27398 | 18 |
| **15** | ARTISTA | 15DPI_1 | day 15 after injury, section 1 | 8996 | 23331 | 18 |
| **16** | ARTISTA | 20DPI_2 | day 20 after injury, section 2 | 11319 | 28987 | 20 |
| **17** | ARTISTA | 30DPI | day 30 after injury | 9252 | 19909 | 17 |
| **18** | ARTISTA | 60DPI | day 60 after injury | 10964 | 20038 | 16 |
| **19** | MOSTA | E9.5_E1S1 | day 9.5 | 5913 | 25568 | 12 |
| **20** | MOSTA | E10.5_E1S1 | day 10.5 | 18408 | 25201 | 13 |
| **21** | MOSTA | E11.5_E1S1 | day 11.5 | 30124 | 26854 | 19 |
| **22** | MOSTA | E12.5_E1S1 | day 12.5 | 51365 | 27810 | 23 |
| **23** | MOSTA | E14.5_E1S1 | day 14.5 | 102519 | 28463 | 26 |
| **24** | MOSTA | E16.5_E1S1 | day 16.5 | 121767 | 28204 | 25 |

Supplementary Table 1 : List of benchmarked Stereo-Seq sections. From left to right, columns represent (i) dataset name, (ii) section name (iii) number of cells (iv) total number of genes before filtering (v) number of ground truth domains.

| | dataset | section | name | n_cells | total genes | ccf | tissue | cell type |
|---|---|---|---|---|---|---|---|---|
| **0** | MBA | C57BL6J_1_053 | section 053 | 34126 | 1120 | 77 | 9 | 19 |
| **1** | MBA | C57BL6J_1_082 | section 082 | 42117 | 1120 | 170 | 10 | 22 |
| **2** | MBA | C57BL6J_1_117 | section 117 | 22862 | 1120 | 59 | 6 | 21 |
| **3** | MBA | C57BL6J_1_111 | section 111 | 35171 | 1120 | 89 | 8 | 20 |
| **4** | MBA | C57BL6J_1_113 | section 113 | 30864 | 1120 | 84 | 8 | 21 |

Supplementary Table 2 : List of benchmarked MERFISH sections. From left to right, columns represent (i) dataset name, (ii) section name (iii) number of cells (iv) total number of genes before filtering (v) number of ground truth domains for the Common Coordinate Framework annotation (vi) number of ground truth domains for the tissue annotation (vii) number of ground truth domains for the cell type annotation.

| original | refined |
|---|---|
| CM | Cardiomyocyte |
| FB | Fibroblast |
| FBmyo | Myofibroblast |
| Macro | Macrophage |
| mo | Myeloid_other |
| DC | Myeloid_other |
| Mast | Myeloid_other |
| Neutrophil | Myeloid_other |
| B | Lymphoid |
| ILC2 | Lymphoid |
| MAIT | Lymphoid |
| NK | Lymphoid |
| T | Lymphoid |
| Treg | Lymphoid |
| Fibrocytes | Fibrocyte |
| Erythrocyte | Other |
| NM | Other |
| Schwann | Other |
| vEC | vEC |
| vEpicardial | vEpicardial |
| vPericyte | vPericyte |
| vSMC | vSMC |
| vSMC1 | vSMC |
| vSMC2 | vSMC |

Supplementary Table 3 : List of correspondances between original cell types provided by the authors and refined cell types used in this study for the mouse cardiac injury dataset.

| original | refined |
|---|---|
| unknown | other |
| Tumor | Tumor cell |
| Plasma_IgG | Plasma IgG |
| CAF | CAF |
| Fibroblasts | Fibroblast |
| T_cell | T cells |
| CAM | CAM |
| Macrophages | Macrophage |
| High_Endothelial_Venules | other |
| M1_macrophages | Macrophage |
| Plasma_IgA | Plasma IgA |
| Germinal_Center_Plasma_IgM_B_cell | Plasma IgM |
| Tumor_Keratin_Pearl | Keratin Pearl |
| Dendritic_cells | other |
| Cytotoxic_IFN_signaling | other |
| Mast_cells | other |
| Cortex_CCL21 | other |

Supplementary Table 4 : List of correspondances between original cell types provided by the authors and refined cell types used in this study for the three dimensional metastatic lymph node.

### Effect of parameter alpha

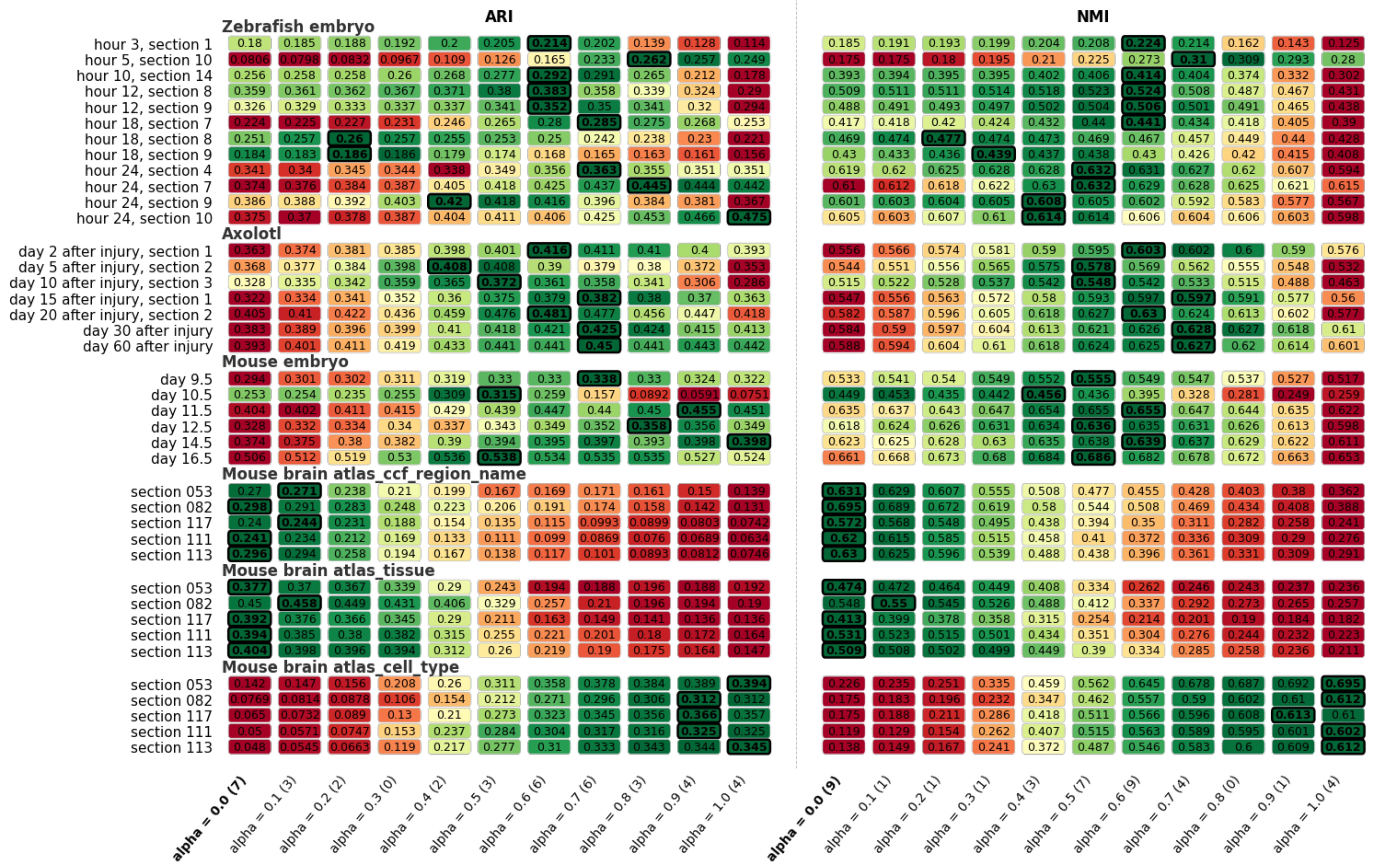


Supplementary figure S1 : Effect of the balance parameter $\alpha$ ($\alpha = \{0, 0.1, 0.2, 0.3, 0.4, 0.5, 0.6, 0.7, 0.8, 0.9, 1\}$) on mean ARI and NMI over 100 seeds for every section and annotation. Kernel parameters fixed as default (sigma_e = 1, snn = 20)

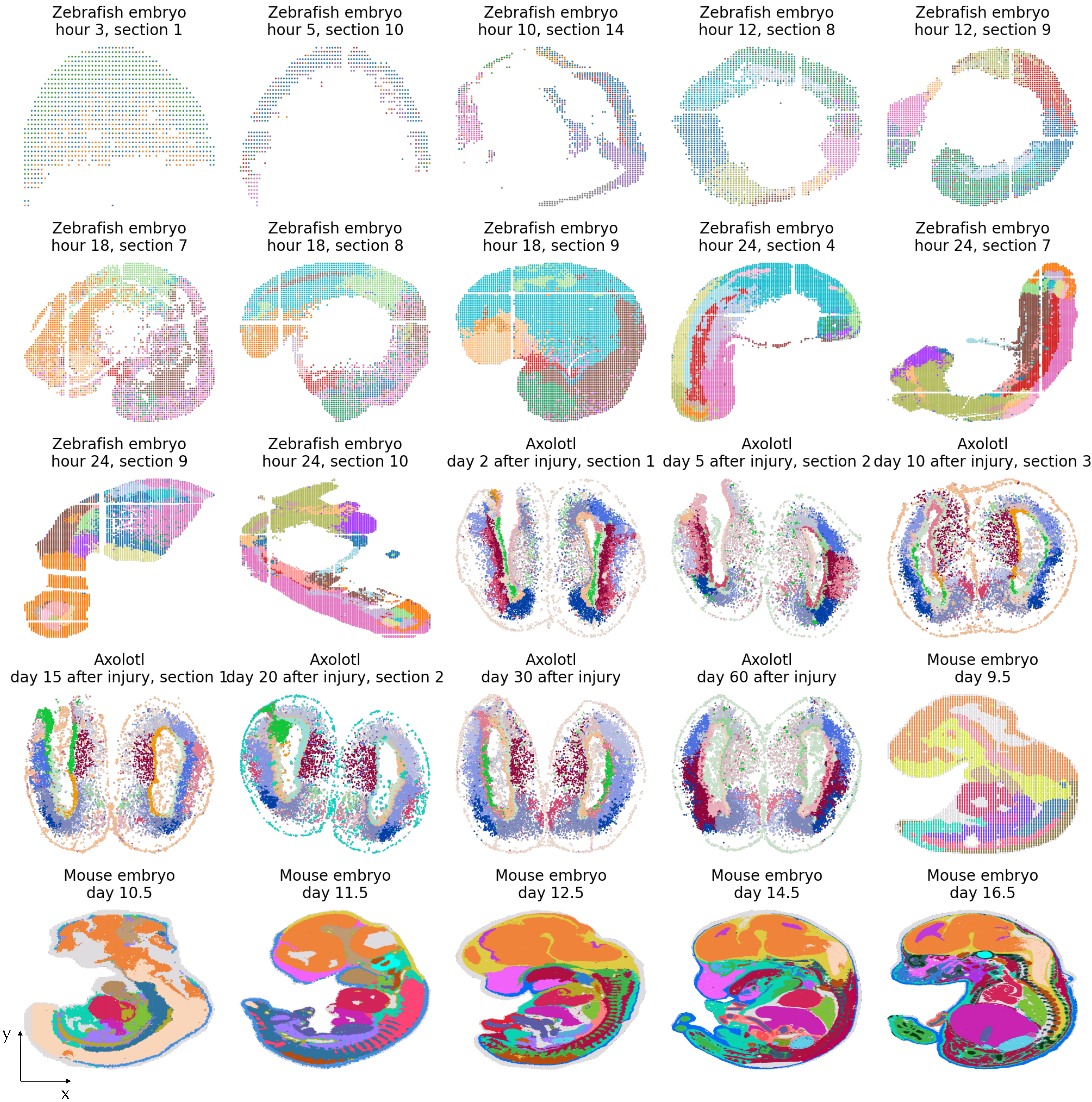


Supplementary figure S2 : Spatial maps of the expert annotations used as ground truth on the 25 Stereo-seq sections used for benchmarking, zebrafish embryo (ZESTA, days 3–24 of development), mouse embryo (MOSTA, days 9.5–16.5) and axolotl telencephalon regeneration (ARTISTA, days 2–60 after injury). Each dot is a cell coloured by its reference domain label.

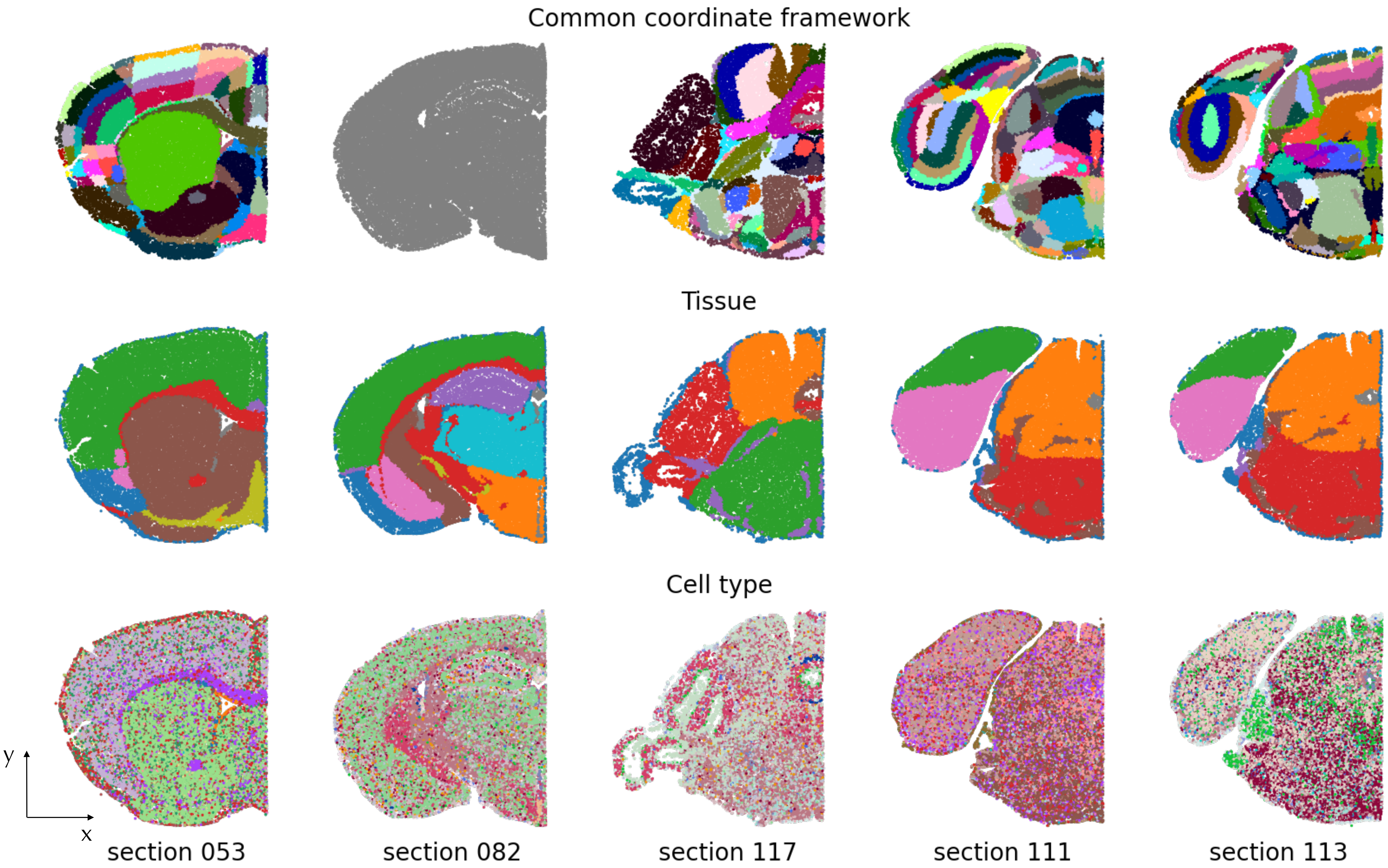


Supplementary figure S3 : Spatial maps of the five coronal sections of the whole-brain MERFISH atlas annotated at three levels of biological granularity: Allen common coordinate framework regions (top row), tissue (middle row) and cell type (bottom row). Each dot is a cell colored by its reference domain label. The grey plot correspond to a sample with 170 ground truth labels that cannot be displayed.

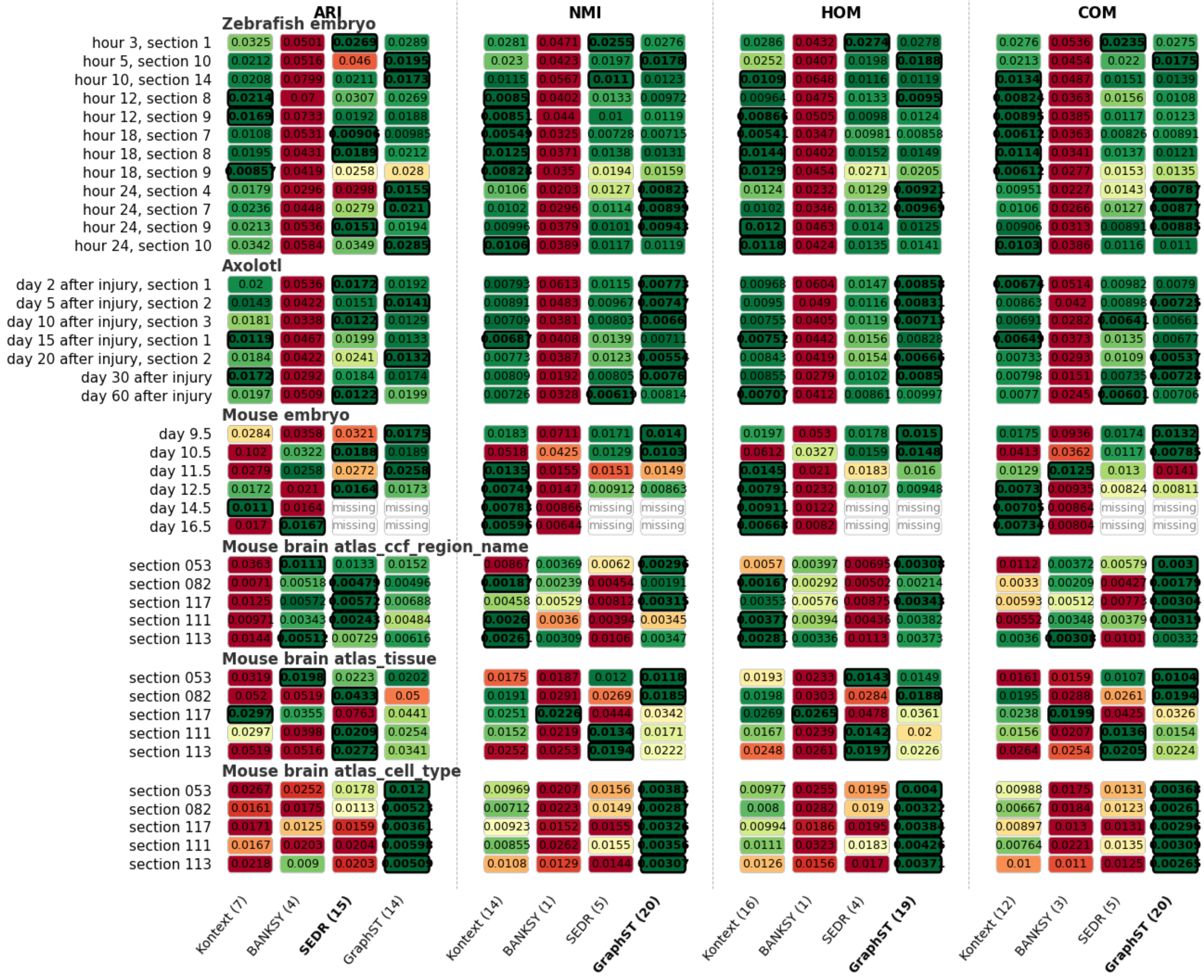


Supplementary figure S4 : Heatmap of the standard deviation of ARI, NMI, homogeneity and completeness computed over 100 random seeds, for every section (rows, grouped by dataset) and every method (columns, grouped by metric). Kontext and BANKSY are deterministic given the data, so the seed only affects the Gaussian-mixture clustering, whereas for SEDR and GraphST the same seed was used for both the embedding and the clustering. Colors are scaled row-wise, with the lowest value outlined in black and reported in bold. The number of sections on which a method is the most stable is given in parentheses in the column labels. Kontext results were obtained with Kontext standard kernel parameters (sigma_e = 1, snn = 20) and varying alpha parameters (stereo-seq : alpha=0.5, MERFISH : ccf :alpha= 0, tissue : alpha=0.1, cell-type alpha=1).

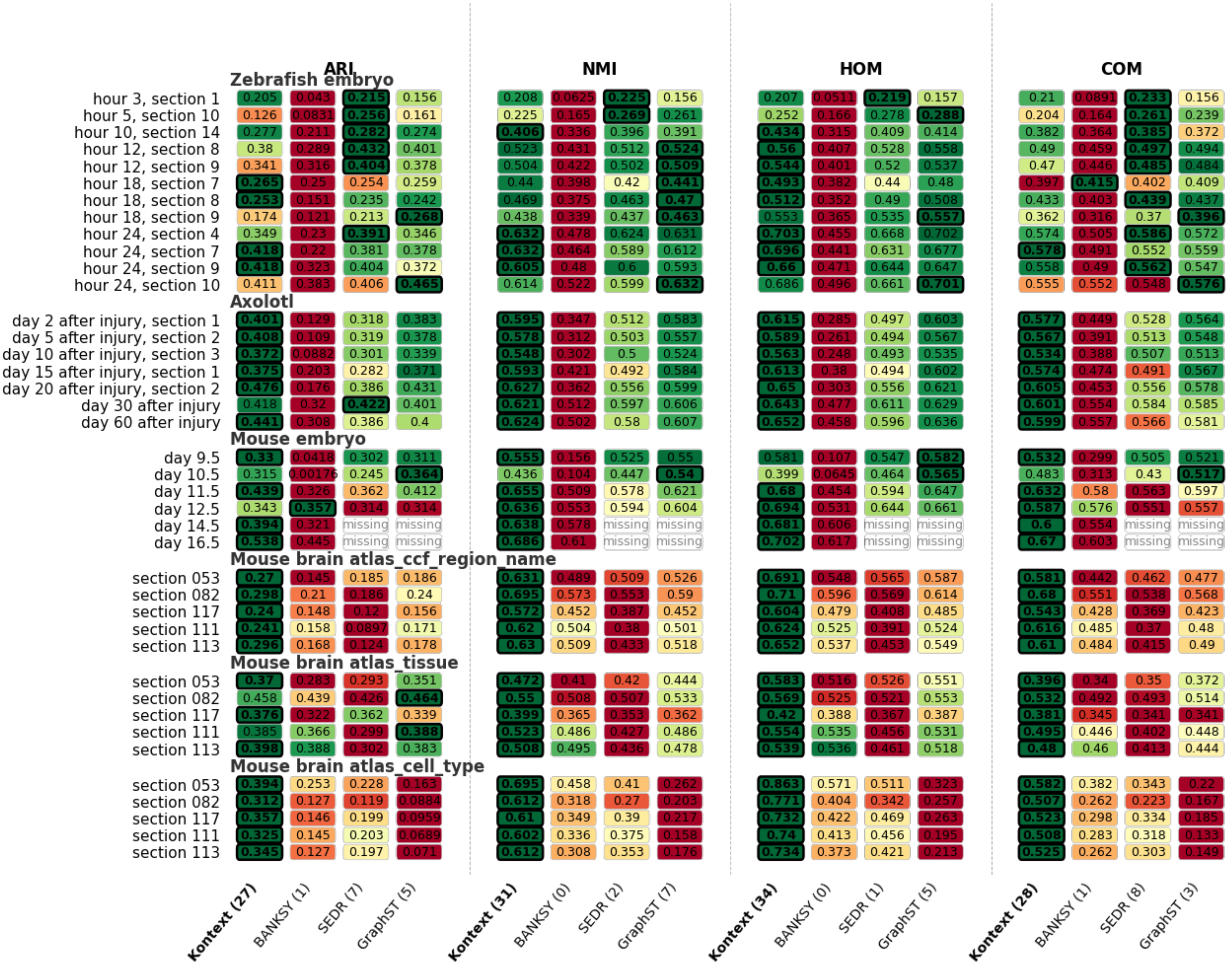


Supplementary figure S5 : Heatmap of the mean ARI, NMI, homogeneity and completeness over 100 random seeds, for every section (rows, grouped by dataset) and every method (columns, grouped by metric). Colors are scaled row-wise from worst (red) to best (green). The best method per section and metric is outlined in black and its value shown in bold. The number of sections won by each method is indicated in parentheses in the column labels. "Missing" entries denote runs that did not complete within the allocated 200 GB of memory. Kontext results were obtained with Kontext standard kernel parameters (sigma_e = 1, snn = 20) and varying alpha parameters (stereo-seq : alpha=0.5, MERFISH : ccf :alpha= 0, tissue : alpha=0.1, cell-type alpha=1).

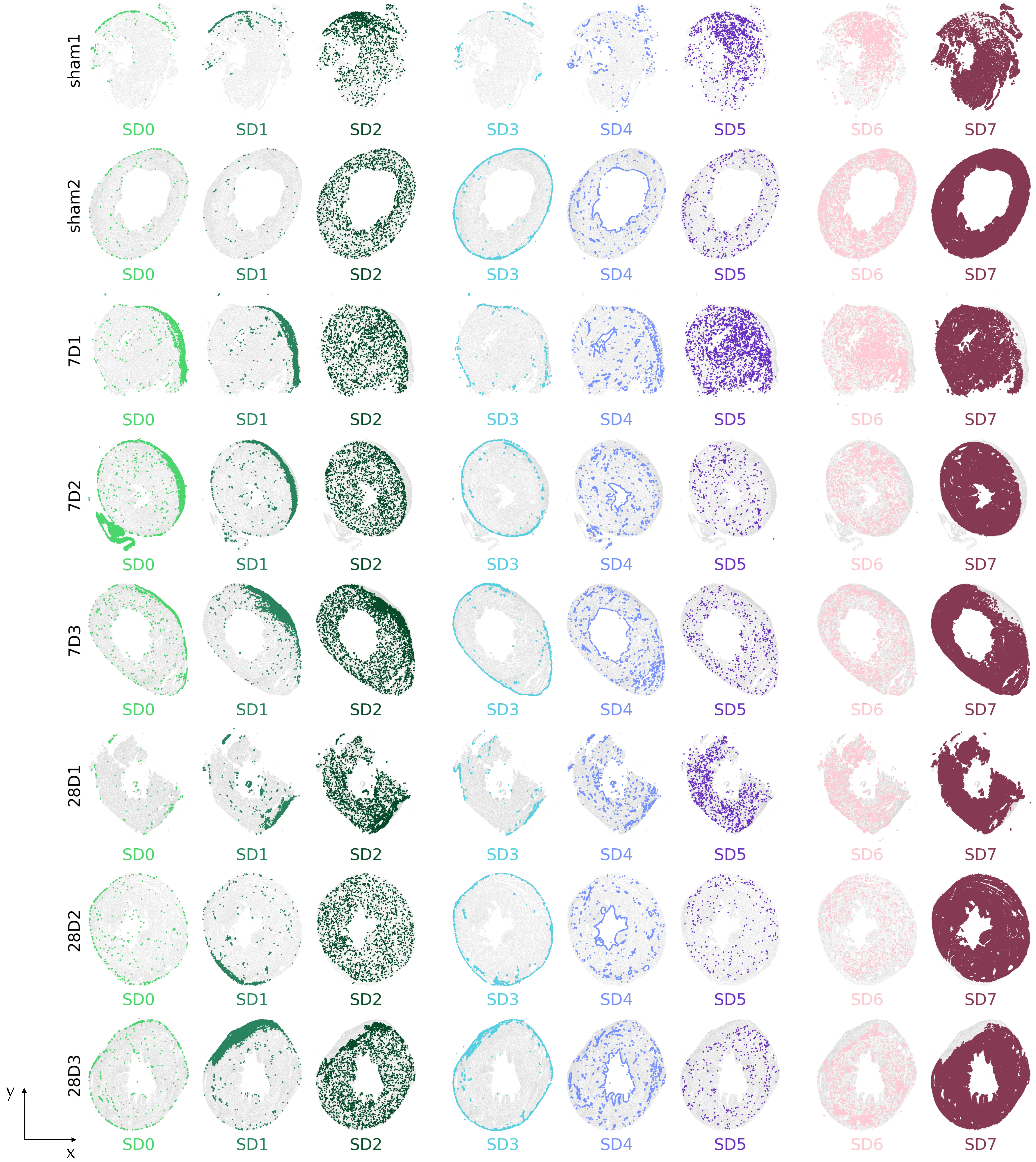


Supplementary figure S6 : Kontext domains in all sections as a grid of spatial coordinates, 8 sections (rows) × 8 domains (columns). In each panel, cells of one domain are highlighted in the domain color over all remaining cells of the section (grey).

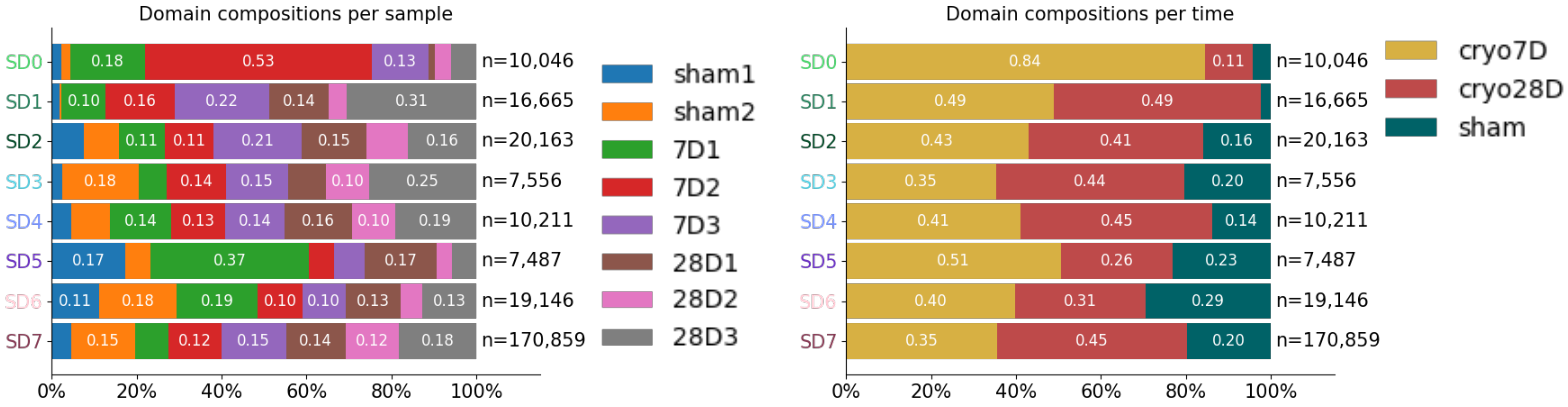


Supplementary figure S7 : Section and temporal composition of the Kontext domains as row-normalized stacked bars giving, for each spatial domain, the proportion of its cells originating from each section (left) and from each time point (right). The total number of cells per domain is indicated on the right.

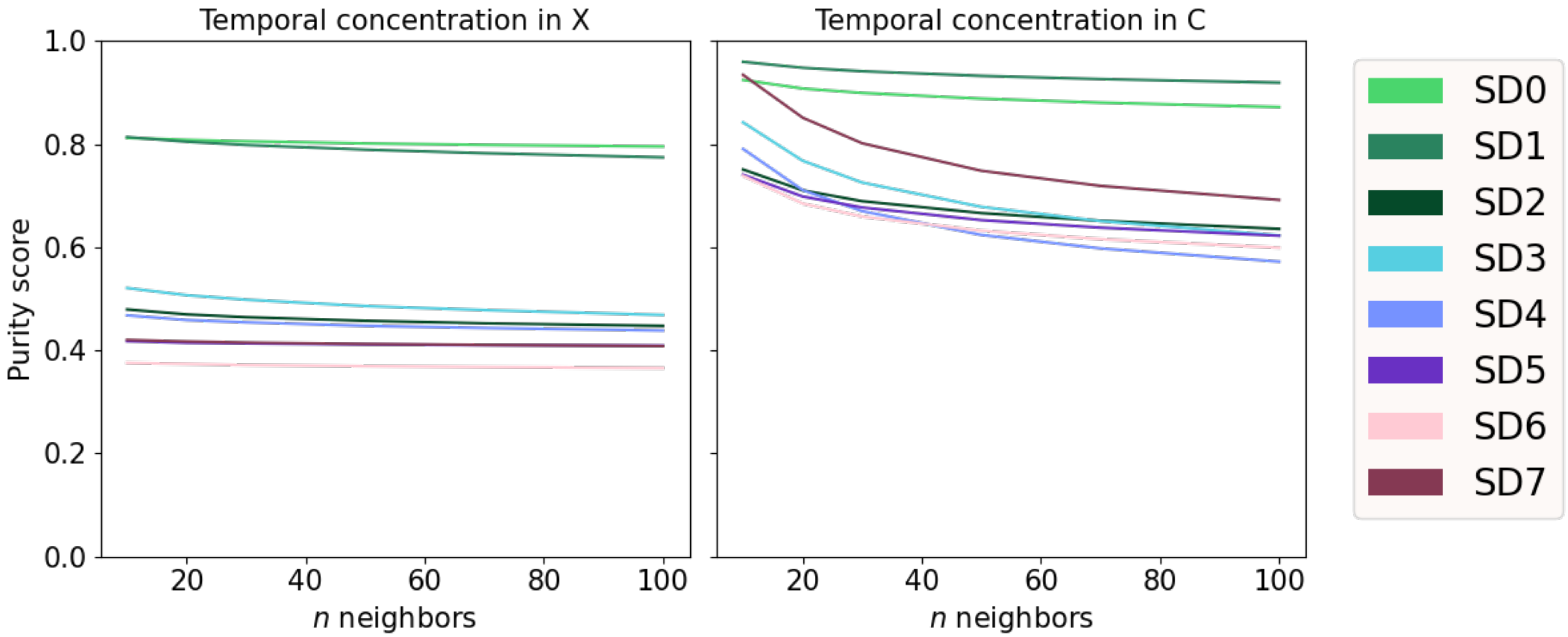


Supplementary figure S8 : Temporal concentration in the expression and microenvironment embedding. For each domain, mean fraction of the k nearest neighbours sharing the same time point (sham, 7D, 28D), computed in the 50-component PCA of the domain-restricted expression matrix X (left) and of the local microenvironment embedding C (right), for k = 10, 20, 30, 50, 70 and 100 neighbours.

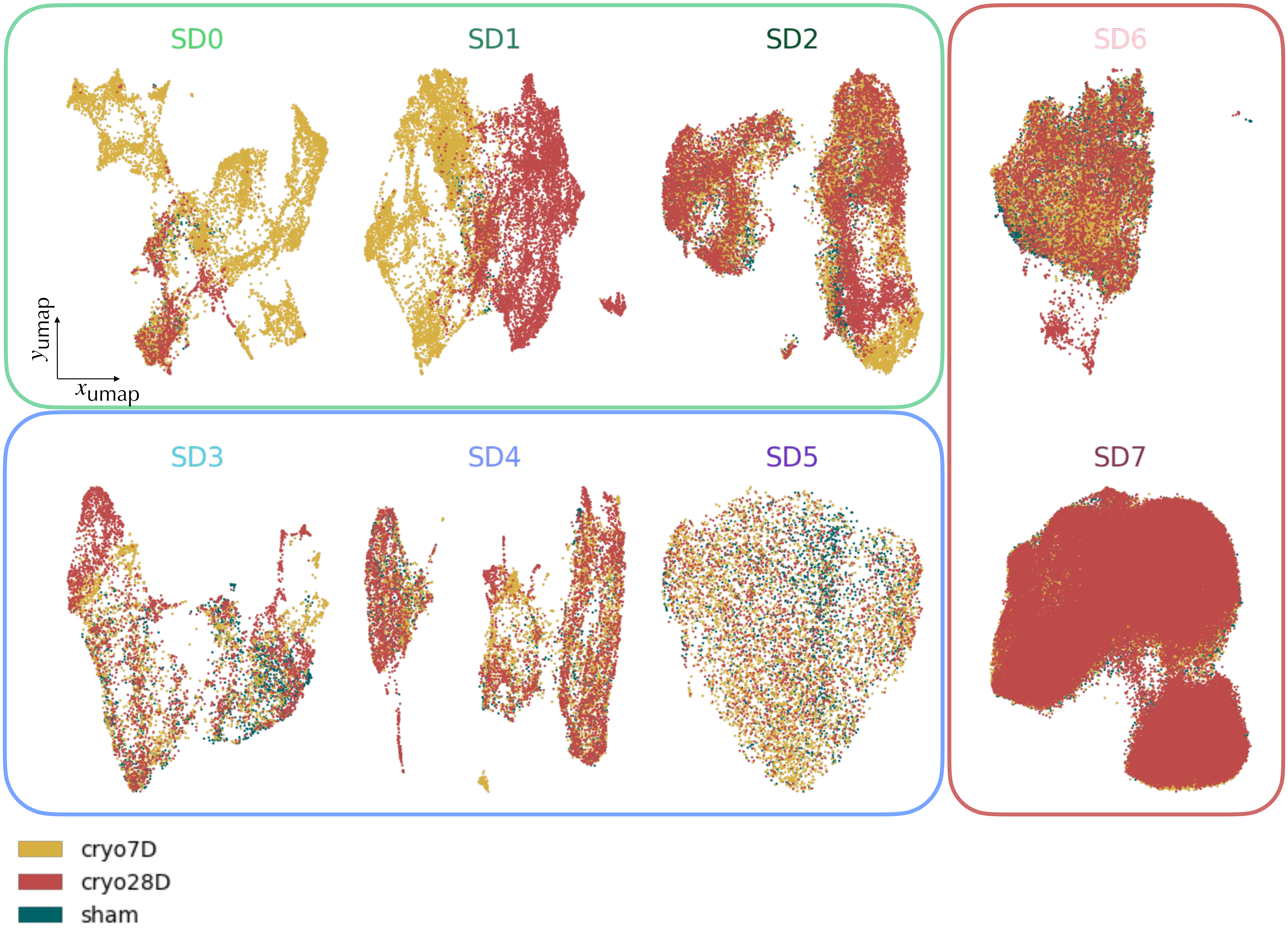


Supplementary figure S9 : 7D and 28D time points are well separated on injury regions SD0 and SD1. Two-dimensional UMAPs of spatially enriched expression embeddings D (50 PCs) computed separately for each domain, with cells colored by time point (sham, 7D, 28D). Panels are grouped as cardiac repair domains (SD0, SD1 and SD2), corronary vascular domains (SD3, SD4 and SD5) and uninjured domains (SD6 and SD7).

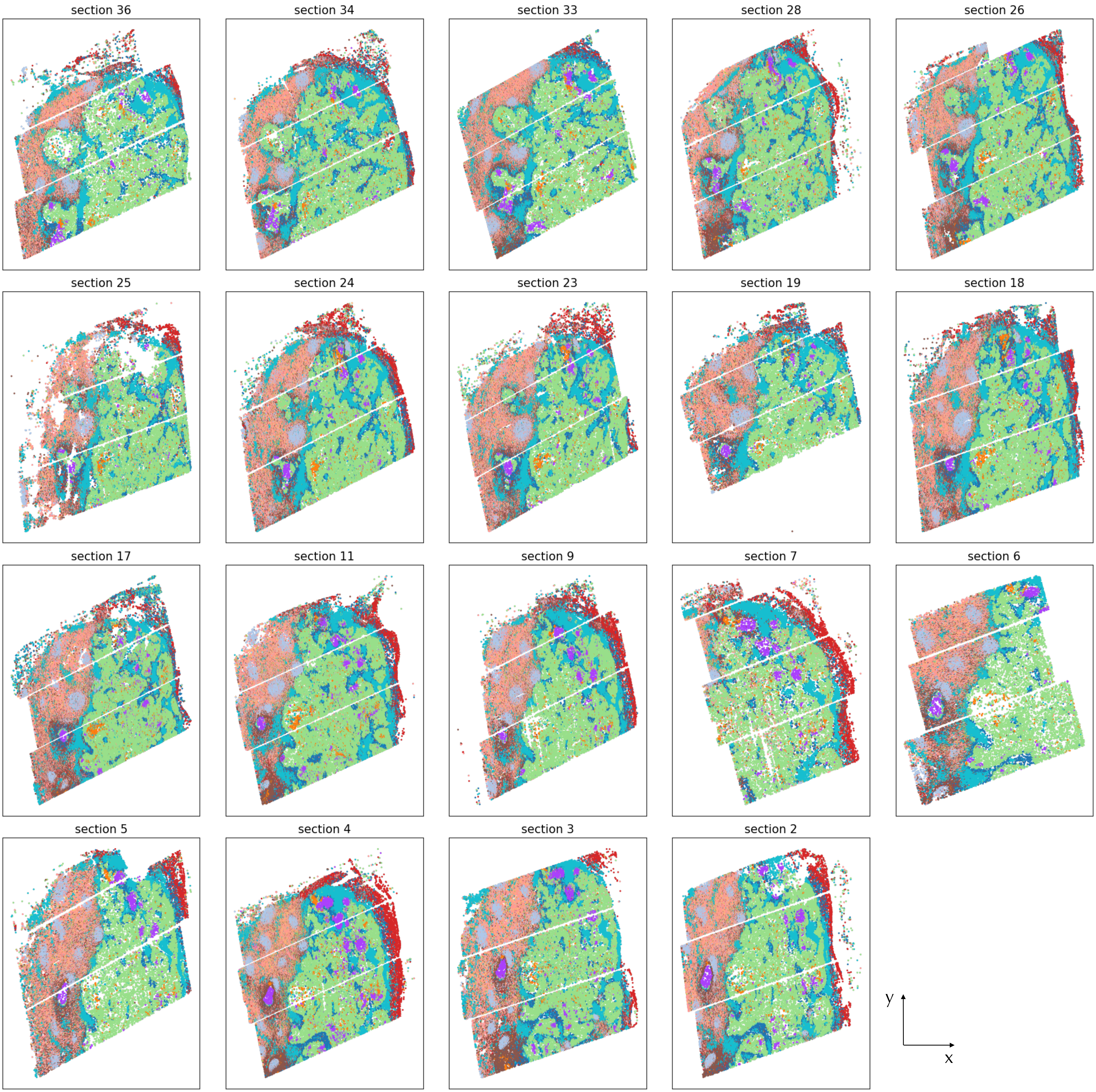


Supplementary figure S10 : Spatial plots of the 19 serial sections (arranged from the top to the bottom section), with cells colored by their refined cell-type annotation.

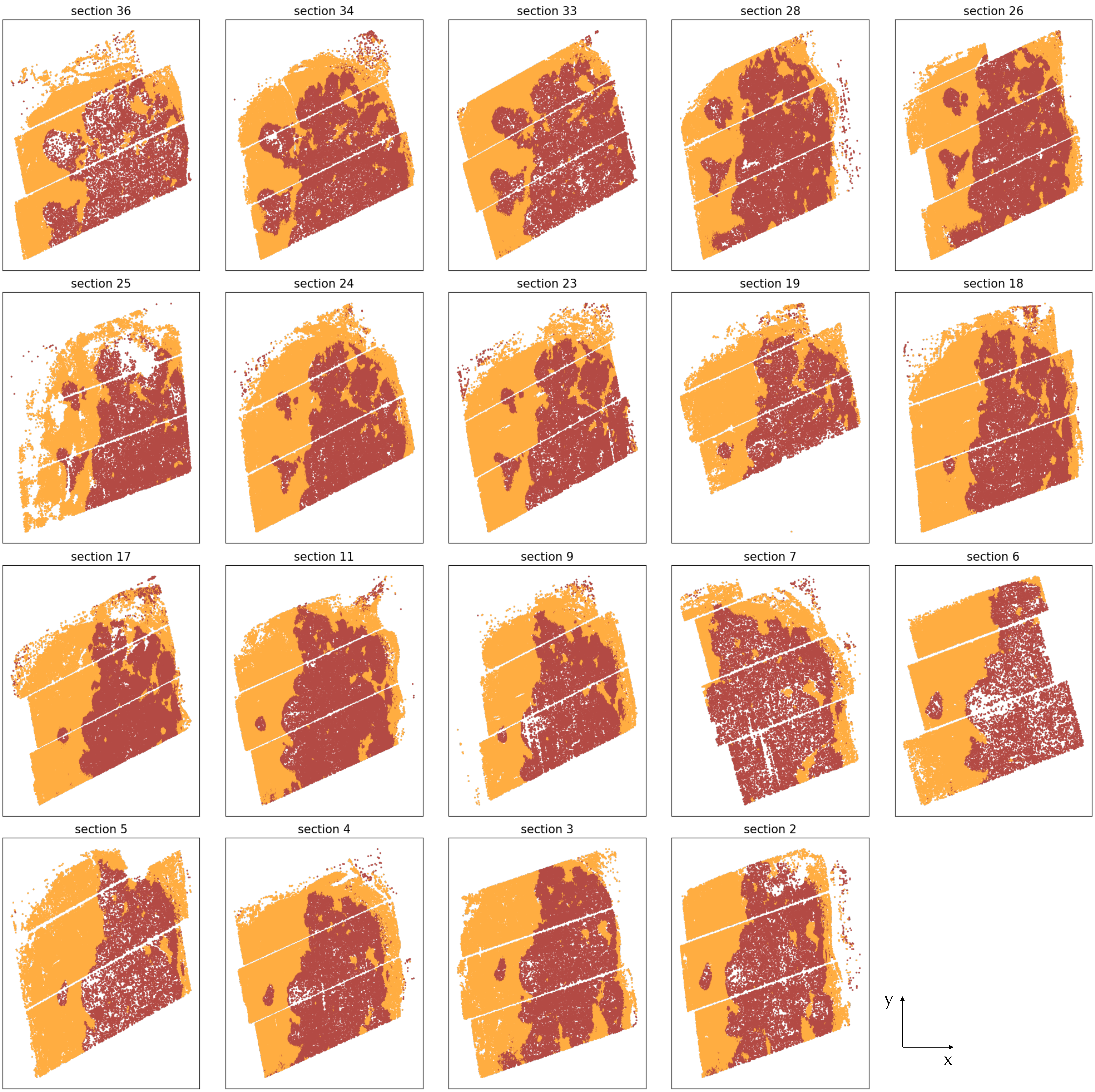


Supplementary figure S11 : Spatial plots of the 19 serial sections (arranged from the top to the bottom section), with cells colored by their Kontext spatial domain (tumor, TME) obtained from the joint three-dimensional analysis.

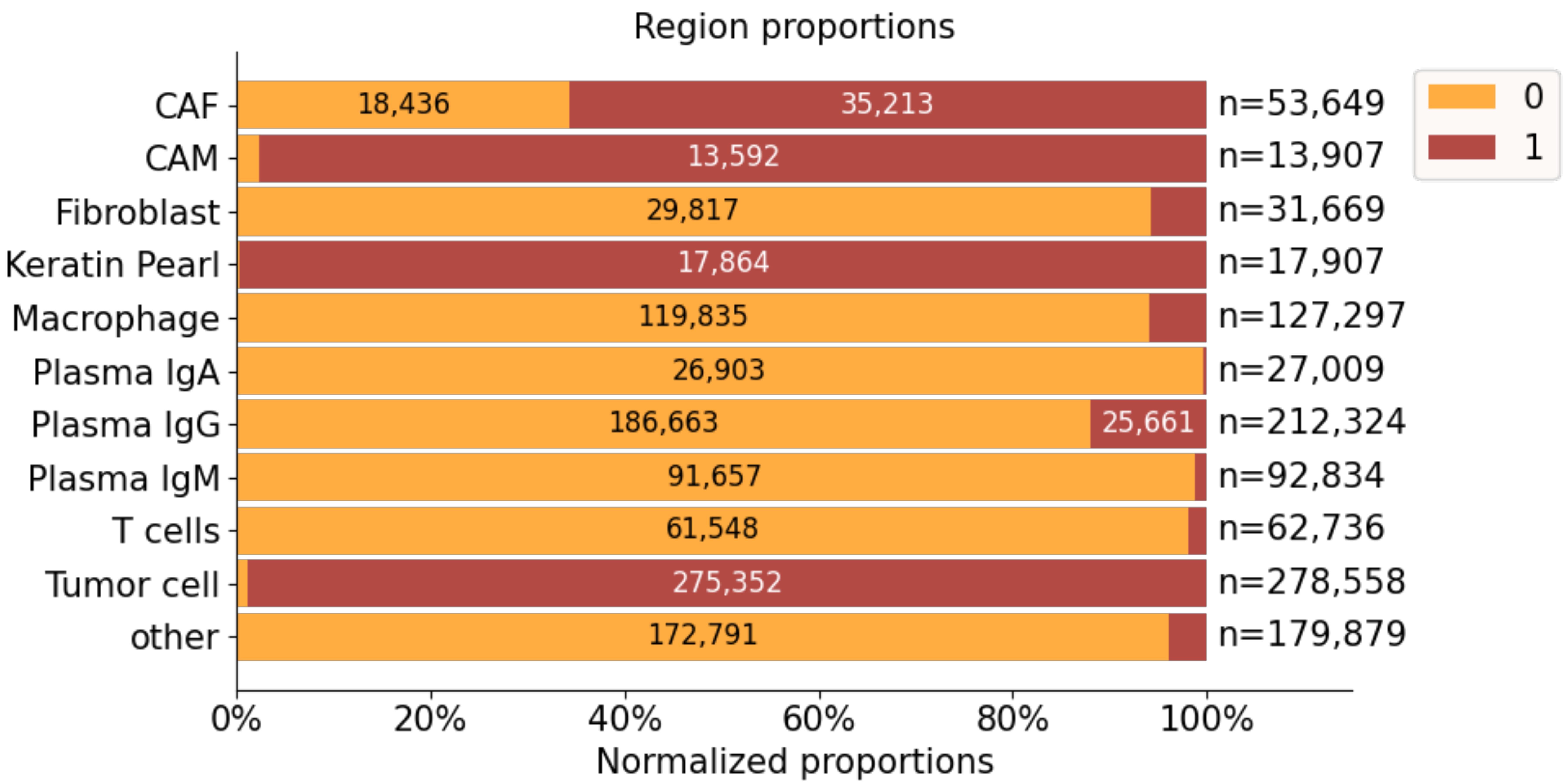


Supplementary figure S12 : Distribution of cell types across Kontext domains as row-normalized stacked bars giving, for each cell type, the proportion of its cells assigned to the tumor and TME domains, pooled over the 19 sections. The total number of cells per cell type is indicated on the right.

3D representation of the tumor region.

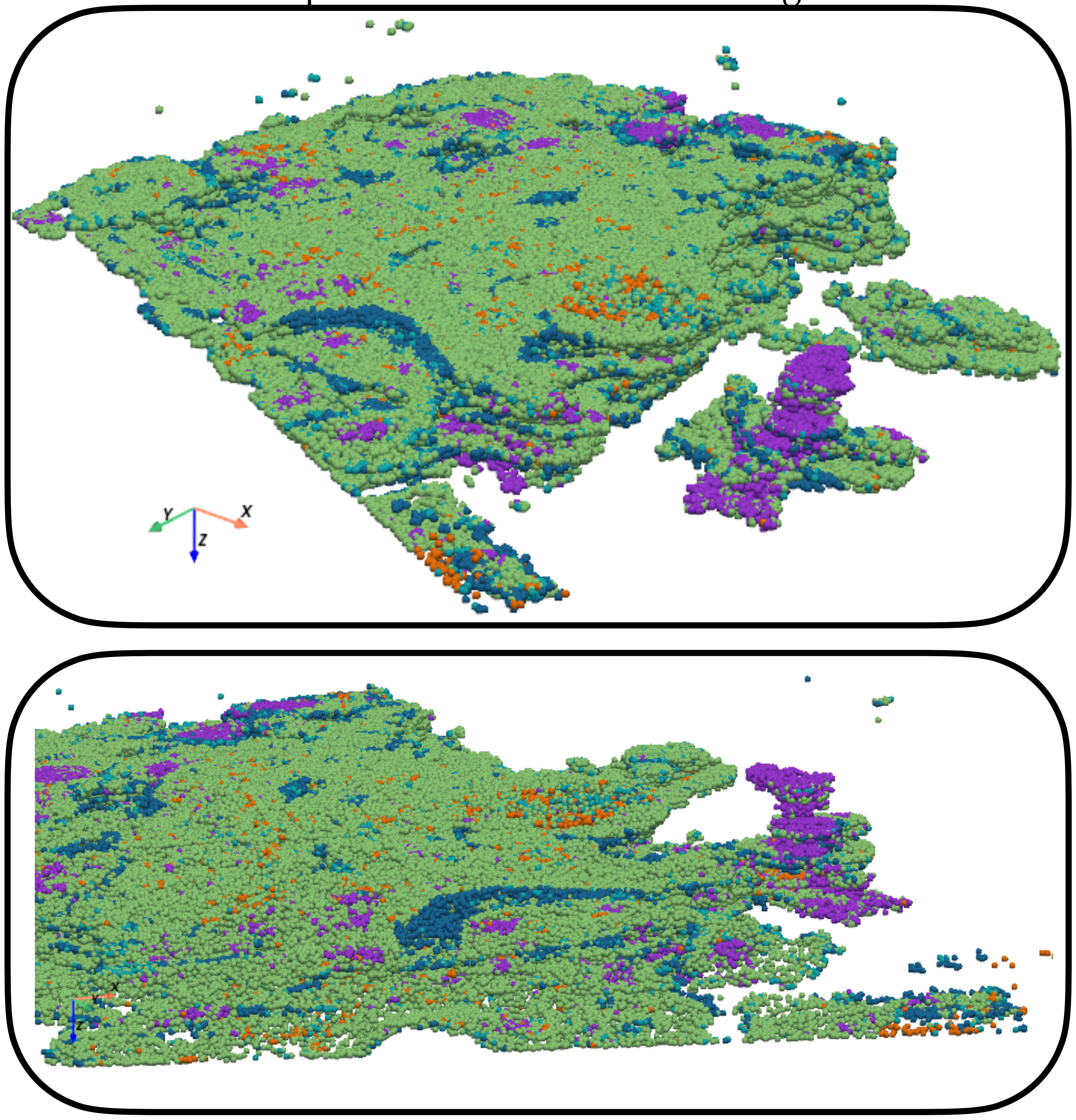


Supplementary figure S13 : Point-cloud renderings of the aligned three-dimensional coordinates, restricted to tumor cells, keratin pearls, CAFs, CAMs and plasma IgG cells that are in the tumor region and colored by cell type. Two camera viewpoints are shown.

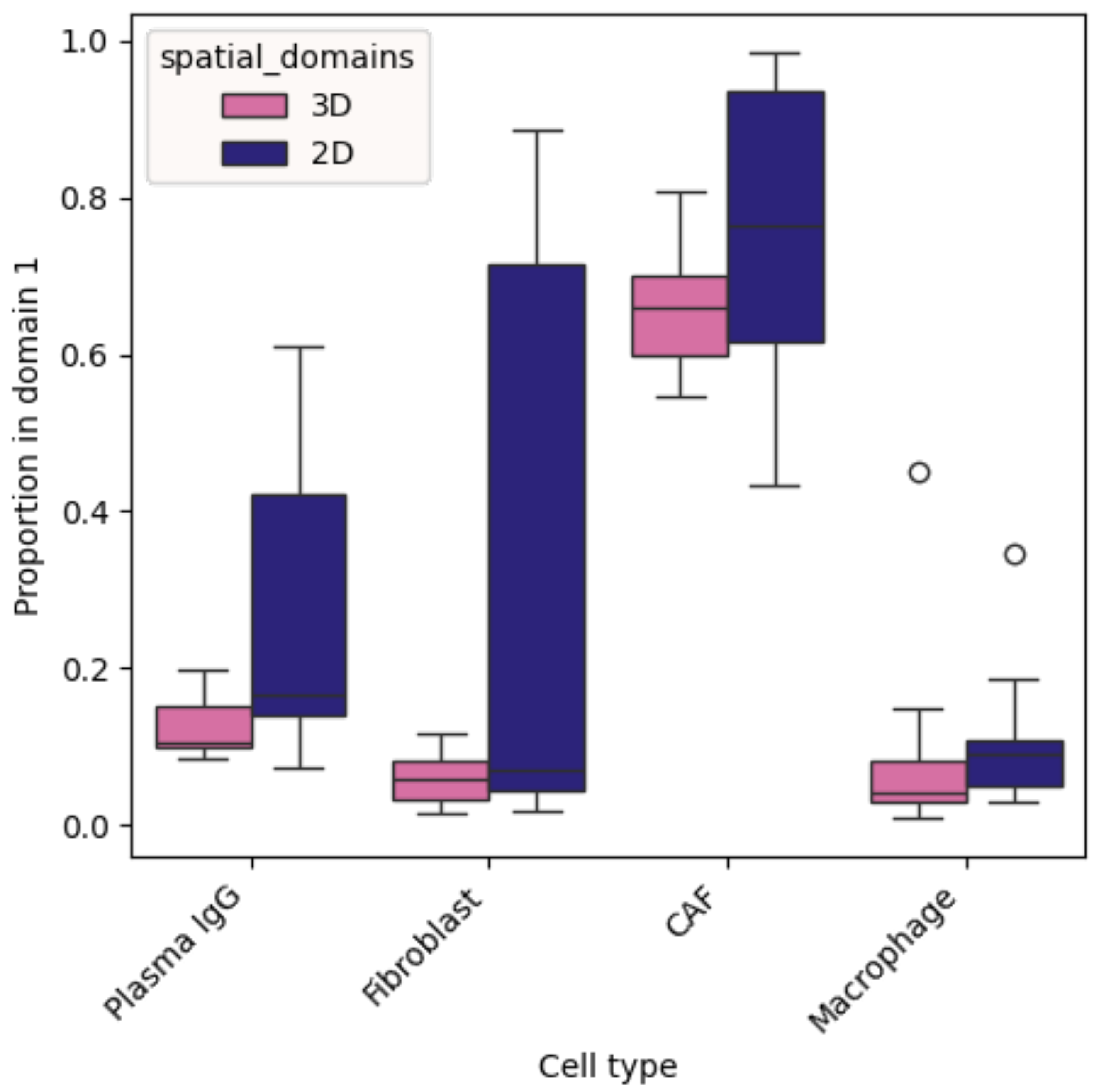


Supplementary figure S14 : Boxplots over the 19 sections of the per-section proportion of cells assigned to the tumor domain, for plasma IgG, fibroblast, CAF and macrophage cells, comparing the domains obtained from the joint three-dimensional analysis (3D) with the domains obtained by running Kontext on each section independently (2D, labels matched to the 3D labels by majority agreement).

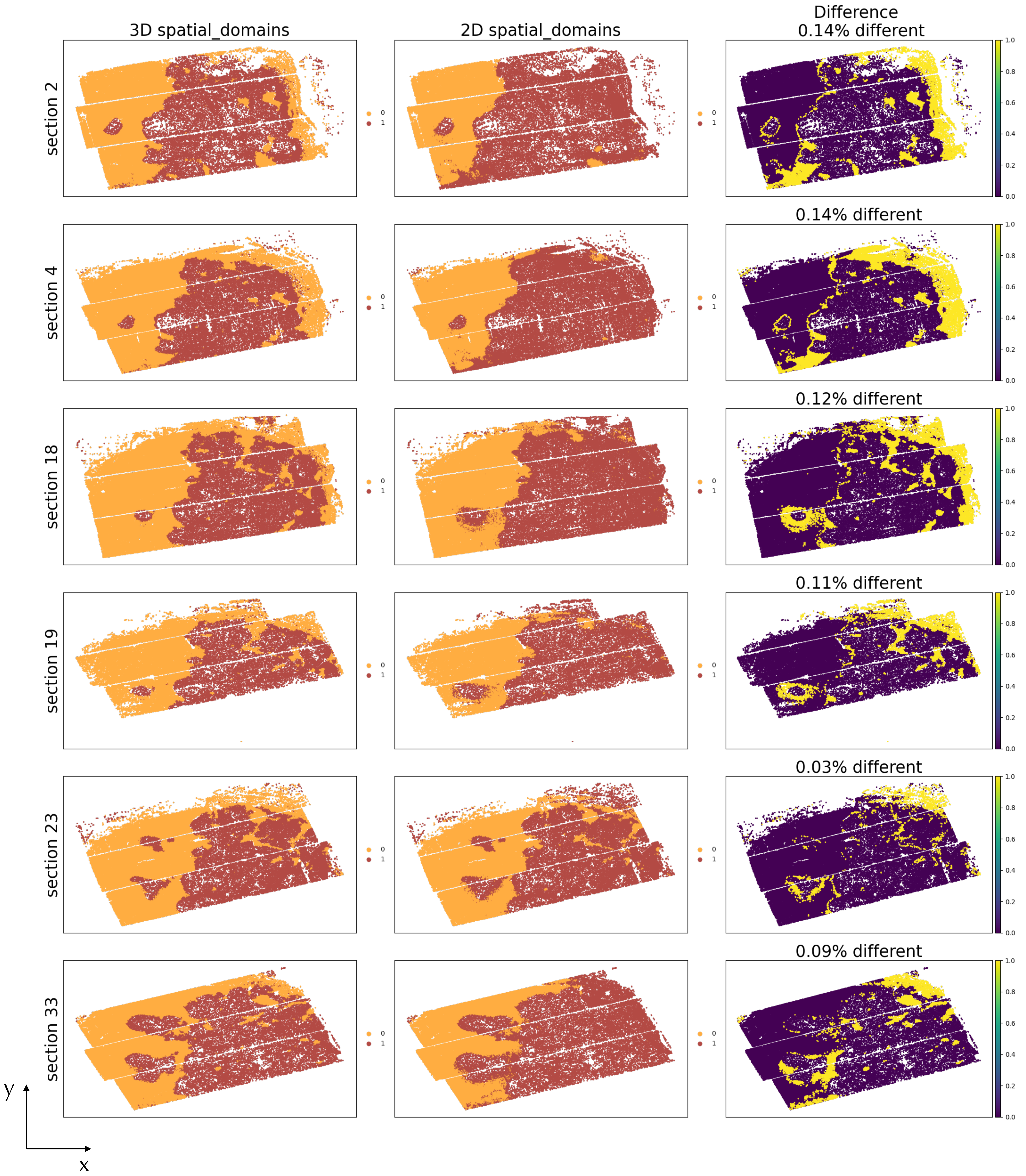


Supplementary figure S15 : Spatial plots of sections 2, 4, 18, 19, 23 and 33 (rows). Left: domains from the joint three-dimensional analysis. Middle: domains obtained by running Kontext on the section alone, with labels matched to the three-dimensional ones. Right: cells whose assignment differs between the two analyses. The number of differing cells out of the total number of cells of the section is given in the panel title.

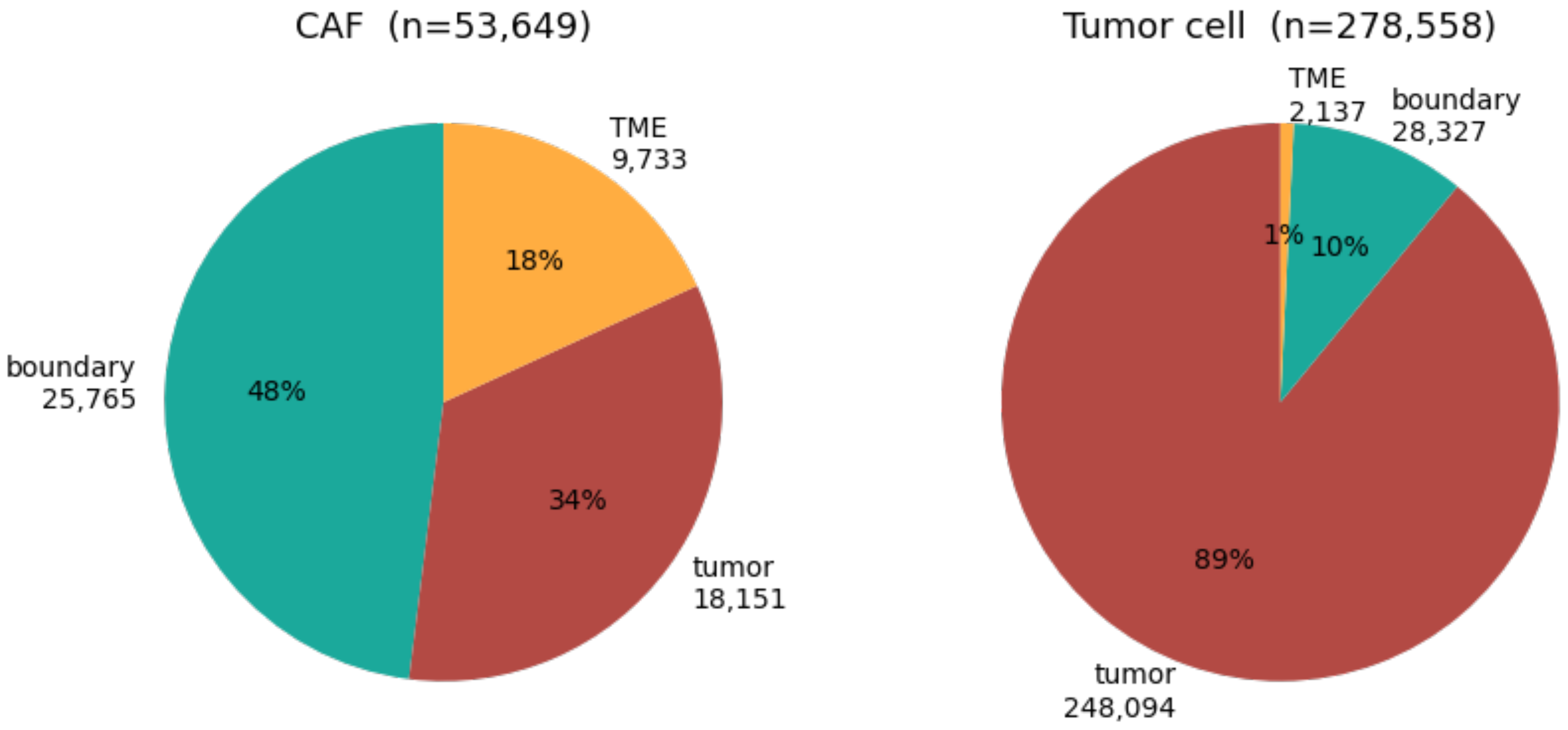


Supplementary figure S16 : Pie plot of the distribution of the CAF cells (left) and tumor cells (right) across the three regions tumor (red), boundary (teal), TME (yellow).

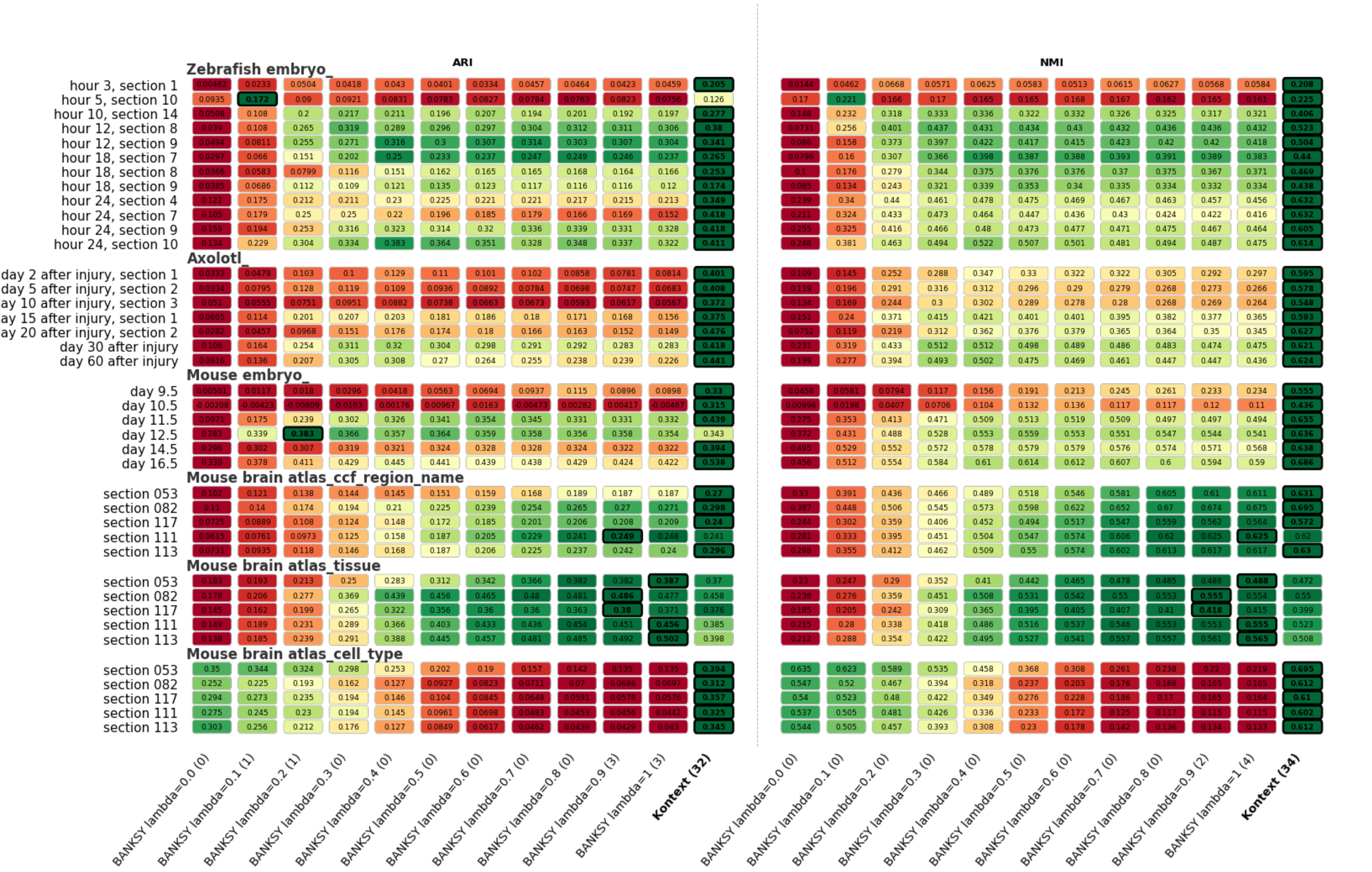


Supplementary figure S17 : Choice of BANKSY's parameter lambda. Heatmap of the mean ARI (left) and NMI (right) over 100 random seeds, for every section (rows, grouped by dataset). Columns represent BANKSY with different values of lambda (from left to right, $\{0, 0.1, 0.2, 0.3, 0.4, 0.5, 0.6, 0.7, 0.8, 0.9, 1\}$. We show Kontext results on the top right column as a reference, with standard parameters $\sigma_e = 1$ and $\ell = 20$, and $\alpha$ set specifically for each dataset (Stereo-seq sections : $\alpha = 0.5$, for MERFISH, ccf : $\alpha = 0$, tissue : $\alpha = 0.1$, cell type : $\alpha = 1$). Colors are scaled row-wise from worst (red) to best (green). The best method per section and metric is outlined in black and its value shown in bold. The number of sections won by each method is indicated in parentheses in the column labels.

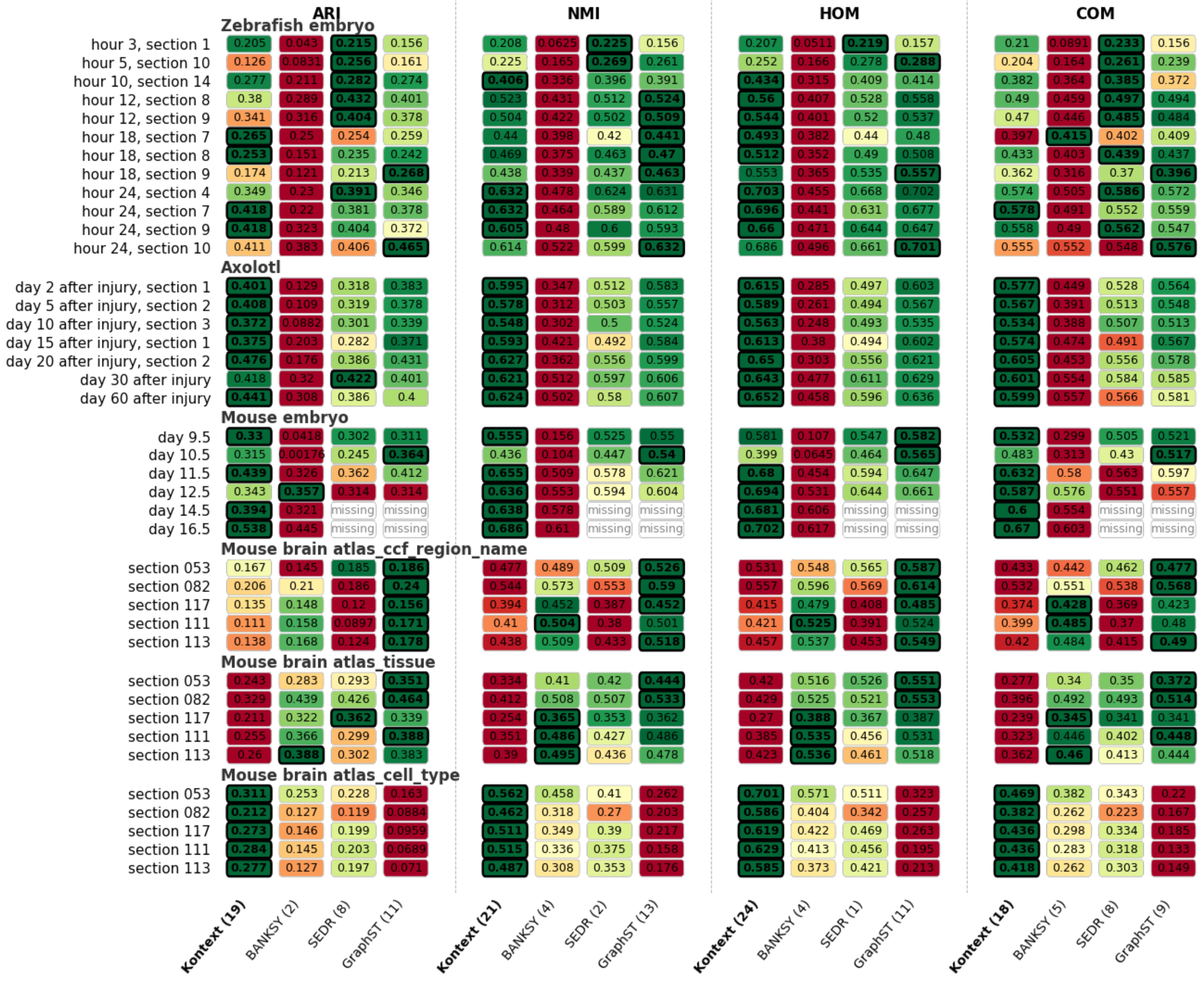


Supplementary figure S18 : Heatmap of the mean ARI, NMI, homogeneity and completeness over 100 random seeds, for every section (rows, grouped by dataset) and every method (columns, grouped by metric). Colors are scaled row-wise from worst (red) to best (green). The best method per section and metric is outlined in black and its value shown in bold. The number of sections won by each method is indicated in parentheses in the column labels. "Missing" entries denote runs that did not complete within the allocated 200 GB of memory. Kontext results were obtained with Kontext standard kernel parameters ($\sigma_e = 1$, $\ell = 20$) and fixed $\alpha = 0.5$.

**ARI**

| | snn=5 (9) | snn=20 (6) | snn=100 (5) | snn=300 (3) | **snn=1000 (11)** | BANKSY (1) | SEDR (2) | GraphST (3) |
|---|---|---|---|---|---|---|---|---|
| **Zebrafish embryo** | | | | | | | | |
| hour 3, section 1 | 0.197 | 0.205 | 0.165 | 0.0386 | 0.0221 | 0.043 | **0.215** | 0.156 |
| hour 5, section 10 | 0.161 | 0.126 | 0.139 | **0.301** | missing | 0.0831 | 0.256 | 0.161 |
| hour 10, section 14 | 0.281 | 0.277 | 0.296 | **0.319** | 0.255 | 0.211 | 0.282 | 0.274 |
| hour 12, section 8 | 0.363 | 0.38 | 0.405 | 0.402 | 0.372 | 0.289 | **0.432** | 0.401 |
| hour 12, section 9 | 0.342 | 0.341 | 0.362 | 0.356 | **0.408** | 0.316 | 0.404 | 0.378 |
| hour 18, section 7 | 0.265 | 0.265 | 0.279 | 0.303 | **0.312** | 0.25 | 0.254 | 0.259 |
| hour 18, section 8 | 0.242 | 0.253 | **0.276** | 0.275 | 0.272 | 0.151 | 0.235 | 0.242 |
| hour 18, section 9 | 0.169 | 0.174 | 0.177 | 0.178 | 0.183 | 0.121 | 0.213 | **0.268** |
| hour 24, section 4 | 0.343 | 0.349 | 0.364 | 0.381 | **0.424** | 0.23 | 0.391 | 0.346 |
| hour 24, section 7 | 0.414 | 0.418 | 0.429 | 0.422 | **0.444** | 0.22 | 0.381 | 0.378 |
| hour 24, section 9 | **0.421** | 0.418 | 0.395 | 0.383 | 0.345 | 0.323 | 0.404 | 0.372 |
| hour 24, section 10 | 0.402 | 0.411 | 0.401 | 0.398 | 0.406 | 0.383 | 0.406 | **0.465** |
| **Axolotl** | | | | | | | | |
| day 2 after injury, section 1 | 0.413 | 0.401 | 0.396 | 0.398 | **0.423** | 0.129 | 0.318 | 0.383 |
| day 5 after injury, section 2 | **0.412** | 0.408 | 0.375 | 0.344 | 0.326 | 0.109 | 0.319 | 0.378 |
| day 10 after injury, section 3 | 0.366 | **0.372** | 0.358 | 0.348 | 0.299 | 0.0882 | 0.301 | 0.339 |
| day 15 after injury, section 1 | **0.377** | 0.375 | 0.36 | 0.349 | 0.352 | 0.203 | 0.282 | 0.371 |
| day 20 after injury, section 2 | 0.472 | **0.476** | 0.452 | 0.43 | 0.435 | 0.176 | 0.386 | 0.431 |
| day 30 after injury | **0.431** | 0.418 | 0.401 | 0.399 | 0.418 | 0.32 | 0.422 | 0.401 |
| day 60 after injury | **0.444** | 0.441 | 0.425 | 0.42 | 0.407 | 0.308 | 0.386 | 0.4 |
| **Mouse embryo** | | | | | | | | |
| day 9.5 | **0.331** | 0.33 | 0.317 | 0.317 | 0.303 | 0.0418 | 0.302 | 0.311 |
| day 10.5 | 0.286 | 0.315 | 0.175 | 0.0742 | 0.0426 | 0.00176 | 0.245 | **0.364** |
| day 11.5 | **0.445** | 0.439 | 0.429 | 0.437 | 0.426 | 0.326 | 0.362 | 0.412 |
| day 12.5 | 0.346 | 0.343 | 0.335 | 0.318 | 0.32 | **0.357** | 0.314 | 0.314 |
| day 14.5 | **0.396** | 0.394 | 0.382 | 0.374 | 0.364 | 0.321 | missing | missing |
| day 16.5 | **0.54** | 0.538 | 0.519 | 0.515 | 0.509 | 0.445 | missing | missing |
| **Mouse brain atlas_ccf_region_name** | | | | | | | | |
| section 053 | 0.137 | **0.27** | 0.174 | 0.172 | 0.164 | 0.145 | 0.185 | 0.186 |
| section 082 | 0.19 | 0.298 | **0.314** | 0.311 | 0.307 | 0.21 | 0.186 | 0.24 |
| section 117 | 0.139 | 0.24 | 0.249 | 0.265 | **0.27** | 0.148 | 0.12 | 0.156 |
| section 111 | 0.126 | 0.241 | 0.278 | **0.283** | 0.27 | 0.158 | 0.0897 | 0.171 |
| section 113 | 0.139 | **0.296** | 0.265 | 0.258 | 0.234 | 0.168 | 0.124 | 0.178 |
| **Mouse brain atlas_tissue** | | | | | | | | |
| section 053 | 0.329 | **0.37** | 0.321 | 0.319 | 0.3 | 0.283 | 0.293 | 0.351 |
| section 082 | 0.434 | 0.458 | **0.497** | 0.49 | 0.468 | 0.439 | 0.426 | 0.464 |
| section 117 | 0.31 | 0.376 | **0.391** | 0.383 | 0.379 | 0.322 | 0.362 | 0.339 |
| section 111 | 0.38 | 0.385 | 0.434 | 0.449 | **0.456** | 0.366 | 0.299 | 0.388 |
| section 113 | 0.406 | 0.398 | **0.457** | 0.441 | 0.417 | 0.388 | 0.302 | 0.383 |
| **Mouse brain atlas_cell_type** | | | | | | | | |
| section 053 | 0.313 | 0.311 | 0.313 | 0.313 | **0.313** | 0.253 | 0.228 | 0.163 |
| section 082 | 0.22 | 0.212 | 0.22 | 0.22 | **0.22** | 0.127 | 0.119 | 0.0884 |
| section 117 | 0.285 | 0.273 | 0.285 | 0.285 | **0.285** | 0.146 | 0.199 | 0.0959 |
| section 111 | 0.26 | **0.284** | 0.26 | 0.26 | 0.26 | 0.145 | 0.203 | 0.0689 |
| section 113 | 0.286 | 0.277 | 0.286 | 0.286 | **0.286** | 0.127 | 0.197 | 0.071 |

**NMI**

| | **snn=5 (10)** | snn=20 (3) | snn=100 (6) | snn=300 (9) | snn=1000 (7) | BANKSY (0) | SEDR (1) | GraphST (4) |
|---|---|---|---|---|---|---|---|---|
| **Zebrafish embryo** | | | | | | | | |
| hour 3, section 1 | 0.208 | 0.208 | 0.186 | 0.0811 | 0.0559 | 0.0625 | **0.225** | 0.156 |
| hour 5, section 10 | 0.254 | 0.225 | 0.24 | **0.289** | missing | 0.165 | 0.269 | 0.261 |
| hour 10, section 14 | 0.406 | 0.406 | **0.41** | 0.4 | 0.345 | 0.336 | 0.396 | 0.391 |
| hour 12, section 8 | 0.503 | 0.523 | 0.539 | **0.542** | 0.511 | 0.431 | 0.512 | 0.524 |
| hour 12, section 9 | 0.501 | 0.504 | **0.515** | 0.507 | 0.514 | 0.422 | 0.502 | 0.509 |
| hour 18, section 7 | 0.438 | 0.44 | 0.441 | 0.436 | 0.435 | 0.398 | 0.42 | **0.441** |
| hour 18, section 8 | 0.462 | 0.469 | **0.487** | 0.477 | 0.468 | 0.375 | 0.463 | 0.47 |
| hour 18, section 9 | 0.432 | 0.438 | 0.441 | 0.443 | 0.437 | 0.339 | 0.437 | **0.463** |
| hour 24, section 4 | 0.628 | 0.632 | **0.635** | 0.63 | 0.621 | 0.478 | 0.624 | 0.631 |
| hour 24, section 7 | 0.628 | 0.632 | **0.633** | 0.63 | 0.622 | 0.464 | 0.589 | 0.612 |
| hour 24, section 9 | **0.607** | 0.605 | 0.595 | 0.579 | 0.552 | 0.48 | 0.6 | 0.593 |
| hour 24, section 10 | 0.606 | 0.614 | 0.612 | 0.609 | 0.608 | 0.522 | 0.599 | **0.632** |
| **Axolotl** | | | | | | | | |
| day 2 after injury, section 1 | **0.605** | 0.595 | 0.581 | 0.569 | 0.574 | 0.347 | 0.512 | 0.583 |
| day 5 after injury, section 2 | **0.585** | 0.578 | 0.555 | 0.535 | 0.516 | 0.312 | 0.503 | 0.557 |
| day 10 after injury, section 3 | **0.549** | 0.548 | 0.534 | 0.521 | 0.489 | 0.302 | 0.5 | 0.524 |
| day 15 after injury, section 1 | **0.597** | 0.593 | 0.579 | 0.564 | 0.558 | 0.421 | 0.492 | 0.584 |
| day 20 after injury, section 2 | **0.629** | 0.627 | 0.612 | 0.601 | 0.591 | 0.362 | 0.556 | 0.599 |
| day 30 after injury | **0.632** | 0.621 | 0.607 | 0.601 | 0.611 | 0.512 | 0.597 | 0.606 |
| day 60 after injury | **0.627** | 0.624 | 0.615 | 0.613 | 0.601 | 0.502 | 0.58 | 0.607 |
| **Mouse embryo** | | | | | | | | |
| day 9.5 | 0.553 | **0.555** | 0.538 | 0.536 | 0.525 | 0.156 | 0.525 | 0.55 |
| day 10.5 | 0.426 | 0.436 | 0.337 | 0.27 | 0.203 | 0.104 | 0.447 | **0.54** |
| day 11.5 | **0.655** | 0.655 | 0.653 | 0.65 | 0.633 | 0.509 | 0.578 | 0.621 |
| day 12.5 | **0.638** | 0.636 | 0.631 | 0.615 | 0.607 | 0.553 | 0.594 | 0.604 |
| day 14.5 | 0.637 | **0.638** | 0.633 | 0.629 | 0.619 | 0.578 | missing | missing |
| day 16.5 | 0.684 | **0.686** | 0.675 | 0.667 | 0.662 | 0.61 | missing | missing |
| **Mouse brain atlas_ccf_region_name** | | | | | | | | |
| section 053 | 0.523 | 0.631 | 0.665 | **0.68** | 0.674 | 0.489 | 0.509 | 0.526 |
| section 082 | 0.573 | 0.695 | 0.745 | **0.751** | 0.747 | 0.573 | 0.553 | 0.59 |
| section 117 | 0.445 | 0.572 | 0.642 | 0.659 | **0.665** | 0.452 | 0.387 | 0.452 |
| section 111 | 0.461 | 0.62 | 0.695 | **0.707** | 0.701 | 0.504 | 0.38 | 0.501 |
| section 113 | 0.487 | 0.63 | 0.683 | **0.688** | 0.676 | 0.509 | 0.433 | 0.518 |
| **Mouse brain atlas_tissue** | | | | | | | | |
| section 053 | 0.451 | 0.472 | 0.483 | **0.494** | 0.482 | 0.41 | 0.42 | 0.444 |
| section 082 | 0.53 | 0.55 | 0.598 | **0.602** | 0.59 | 0.508 | 0.507 | 0.533 |
| section 117 | 0.342 | 0.399 | 0.452 | 0.453 | **0.463** | 0.365 | 0.353 | 0.362 |
| section 111 | 0.479 | 0.523 | 0.565 | 0.594 | **0.597** | 0.486 | 0.427 | 0.486 |
| section 113 | 0.485 | 0.508 | 0.568 | **0.583** | 0.571 | 0.495 | 0.436 | 0.478 |
| **Mouse brain atlas_cell_type** | | | | | | | | |
| section 053 | 0.637 | 0.562 | 0.637 | 0.637 | **0.637** | 0.458 | 0.41 | 0.262 |
| section 082 | 0.532 | 0.462 | 0.532 | 0.532 | **0.532** | 0.318 | 0.27 | 0.203 |
| section 117 | 0.563 | 0.511 | 0.563 | 0.563 | **0.563** | 0.349 | 0.39 | 0.217 |
| section 111 | 0.537 | 0.515 | **0.537** | 0.537 | 0.537 | 0.336 | 0.375 | 0.158 |
| section 113 | 0.565 | 0.487 | 0.565 | 0.565 | **0.565** | 0.308 | 0.353 | 0.176 |

Supplementary figure S19 : Effect of parameter $\ell$ (called snn in Kontext package). Heatmap of the mean ARI (left), NMI (right) over 100 random seeds, for every section (rows, grouped by dataset). From left to right, columns correspond to Kontext with increasing values of $\ell$ in $\{5,20,100,300,1000\}$, then BANKSY, SEDR and GraphST as references. Colors are scaled row-wise from worst (red) to best (green). The best method per section and metric is outlined in black and its value shown in bold. The number of sections won by each method is indicated in parentheses in the column labels. "Missing" entries denote runs that did not complete within the allocated 200 GB of memory, or when the number of neighbors was higher than the number of observations in the section. Kontext results were obtained with Kontext standard kernel parameter $\sigma_e = 1$ and varying $\alpha$ (Stereo-seq sections : $\alpha = 0.5$, for MERFISH, ccf : $\alpha = 0$, tissue : $\alpha = 0.1$, cell type : $\alpha = 1$).

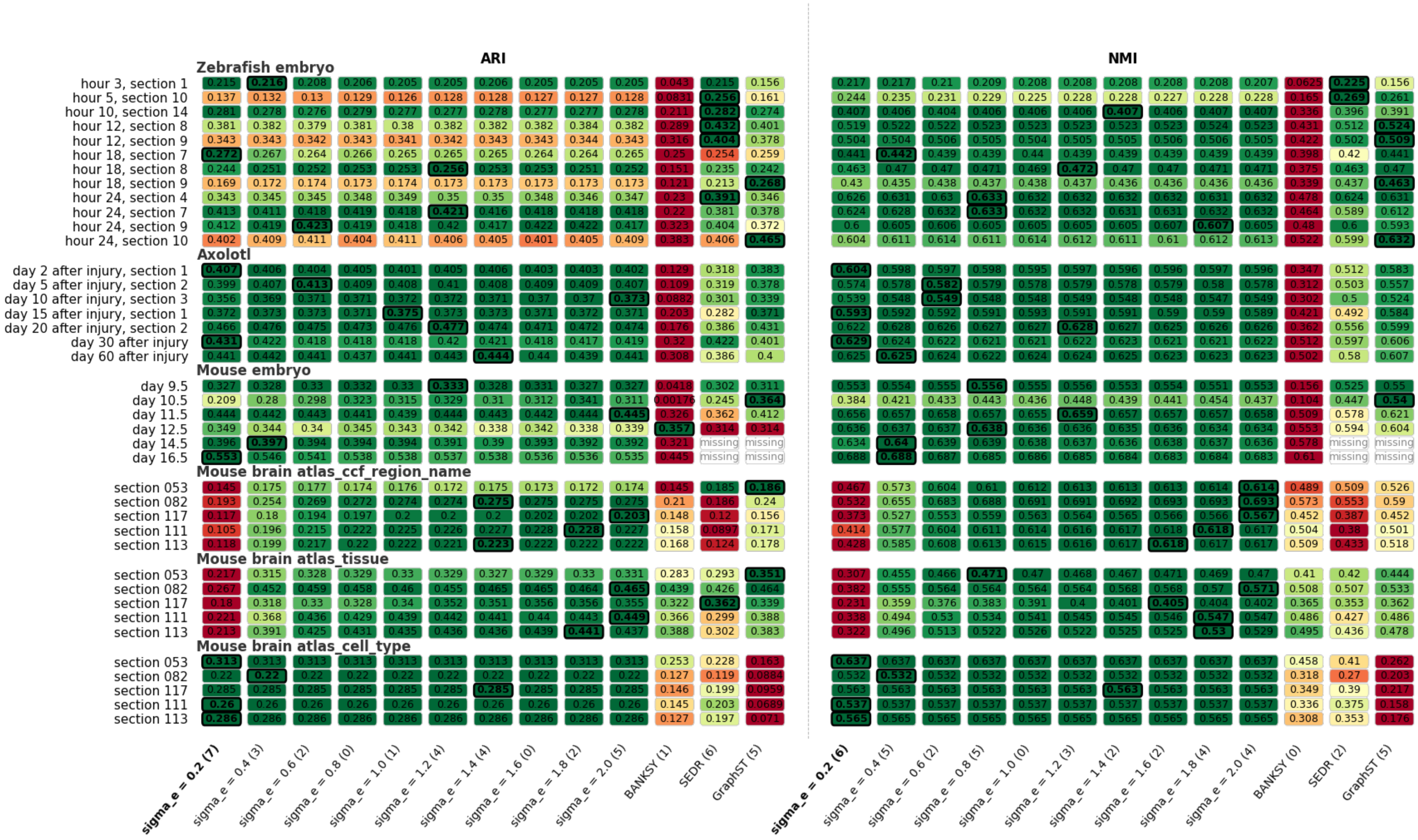


Supplementary figure S20 : Effect of parameter $\sigma_e$ (called sigma_e in Kontext package). Heatmap of the mean ARI (left), NMI (right) over 100 random seeds, for every section (rows, grouped by dataset). From left to right, columns correspond to Kontext with increasing values of $\sigma_e$ in $\{0.2, 0.4, 0.6, 0.8, 1, 1.2, 1.4, 1.6, 1.8, 2\}$, then BANKSY, SEDR and GraphST as references. Colors are scaled row-wise from worst (red) to best (green). The best method per section and metric is outlined in black and its value shown in bold. The number of sections won by each method is indicated in parentheses in the column labels. "Missing" entries denote runs that did not complete within the allocated 200 GB of memory, or when the number of neighbors was higher than the number of observations in the section. Kontext results were obtained with Kontext standard kernel parameter $\ell = 20$ and varying $\alpha$ (Stereo-seq sections : $\alpha = 0.5$, for MERFISH, ccf : $\alpha = 0$, tissue : $\alpha = 0.1$, cell type : $\alpha = 1$).

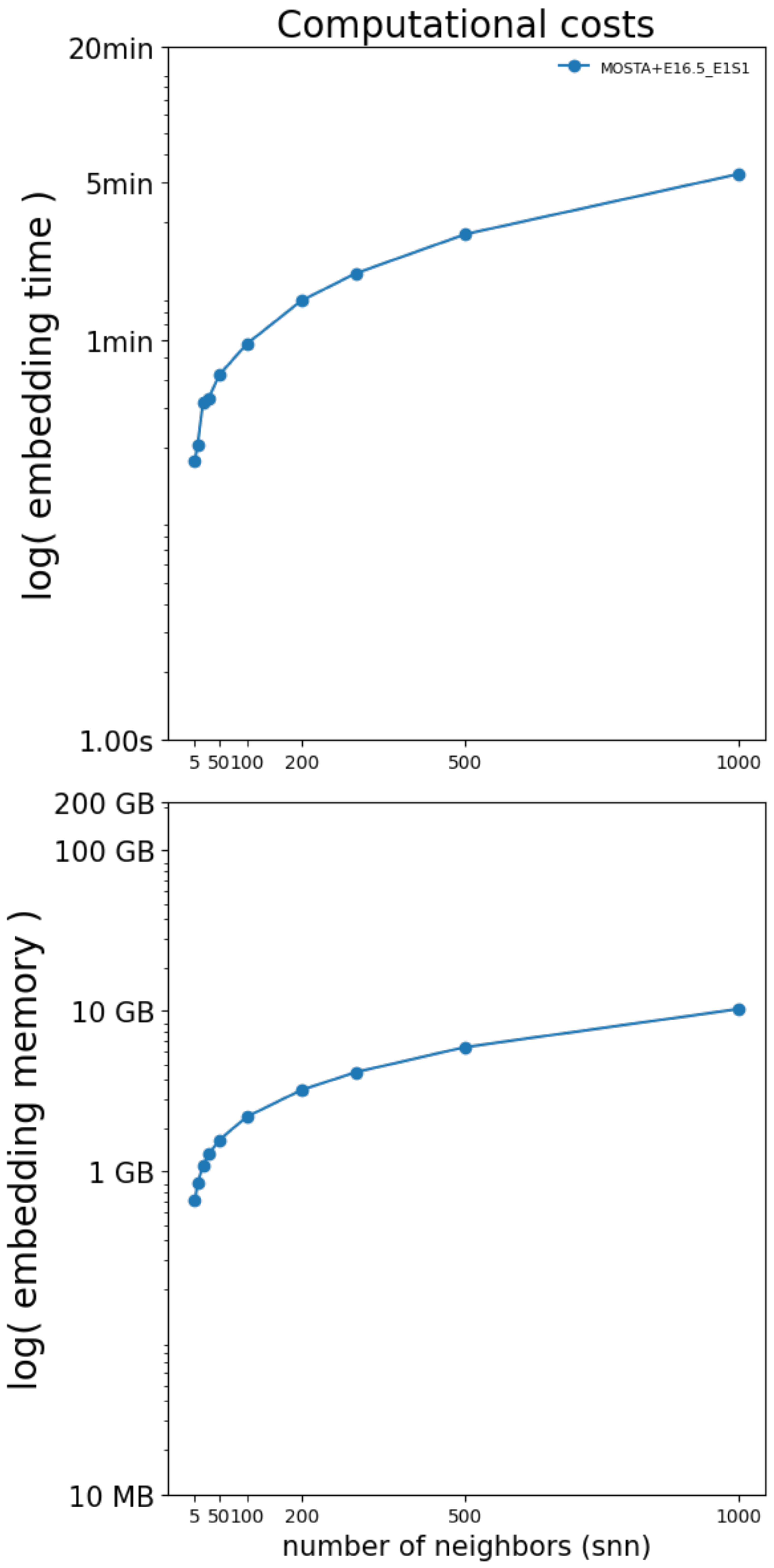


Supplementary figure S21 : Computational cost of the embedding step as a function of the value of parameter $\ell$ (called snn in Kontext package), for the largest benchmarked section E18.5_E1S1 from MOSTA dataset, that contains 121767 cells. Wall-clock time (top) and peak resident memory (bottom), both on a logarithmic scale, with points showing the mean over 100 runs for each value of $\ell$ (snn).

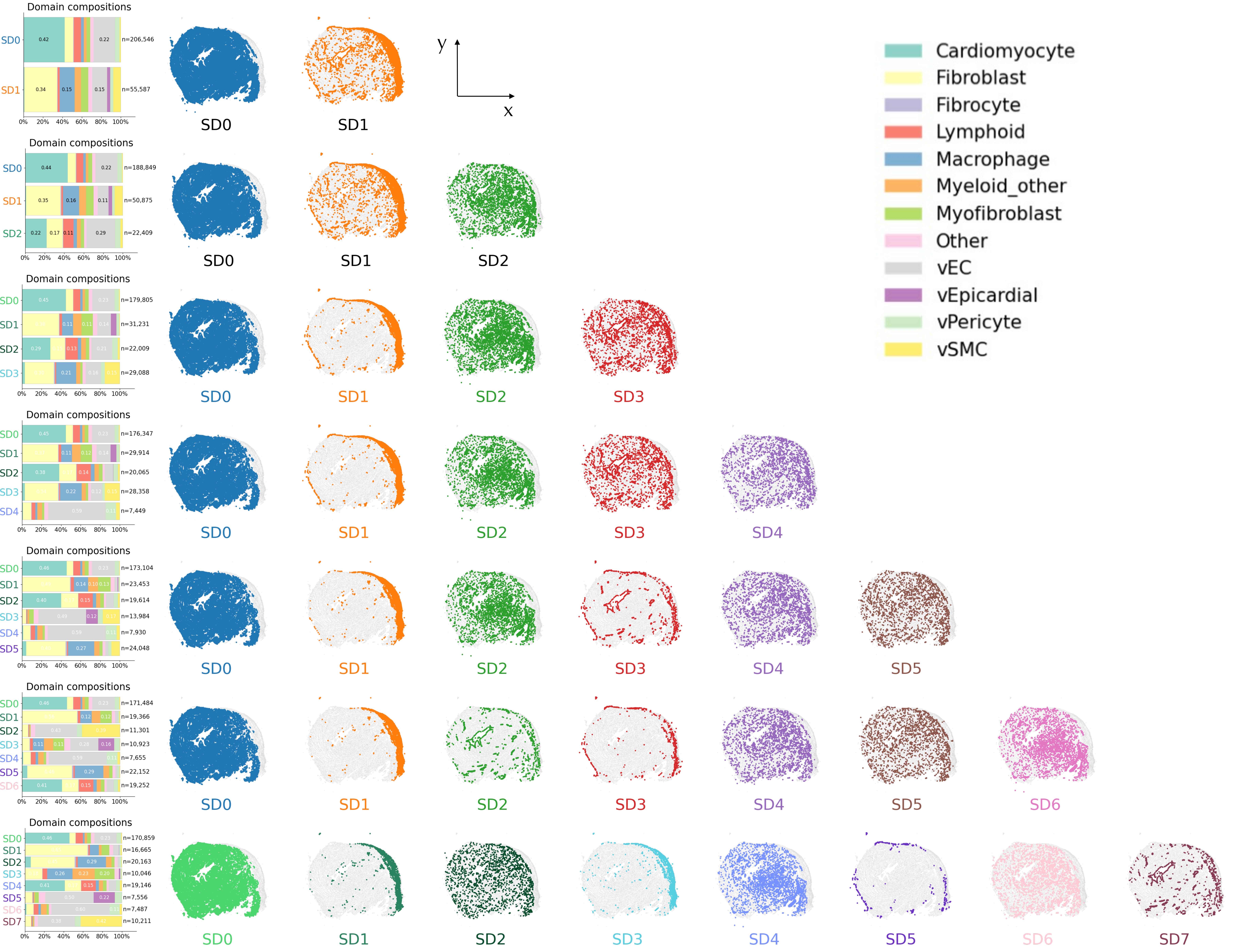


Supplementary figure S22 : Each row correspond to the spatial domains obtained with Kontext for various values of $N_D$ in $\{2,3,4,5,6,7\}$ and fixed set of parameters ($\sigma_e = 1$, $\ell = 20$, $\alpha = 0.5$). Left column : cell type proportions, other columns : spatial display of each spatial domain separately for sample SD1.
